# Modelling biochemical gradients in vitro to control cell compartmentalization in a microengineered 3D model of the intestinal epithelium

**DOI:** 10.1101/2021.12.13.472418

**Authors:** Gizem Altay, Aina Abad-Lázaro, Emilio J. Gualda, Jordi Folch, Claudia Insa, Sébastien Tosi, Xavier Hernando-Momblona, Eduard Batlle, Pablo Loza-Álvarez, Vanesa Fernández-Majada, Elena Martinez

## Abstract

Gradients of signaling pathways within the intestinal stem cell (ISC) niche are instrumental for cellular compartmentalization and tissue function, yet how are they sensed by the epithelium is still not fully understood. Here we present a new in vitro model of the small intestine based on primary epithelial cells (i), apically accessible (ii), with native tissue mechanical properties and controlled mesh size (iii), 3D villus-like architecture (iv), and precisely controlled biomolecular gradients of the ISC niche (v). Biochemical gradients are formed through hydrogel-based scaffolds by free diffusion from a source to a sink chamber. To confirm the establishment of spatiotemporally controlled gradients, we employ light-sheet fluorescence microscopy and in-silico modelling. The ISC niche biochemical gradients coming from the stroma and applied along the villus axis lead to the in vivo-like compartmentalization of the proliferative and differentiated cells, while changing the composition and concentration of the biochemical factors affects the cellular organization along the villus axis. This novel 3D in vitro intestinal model derived from organoids recapitulates both the villus-like architecture and the gradients of ISC biochemical factors, thus opening the possibility to study in vitro the nature of such gradients and the resulting cellular response.

## 1. Introduction

The small intestine is lined by a monolayer of epithelial cells organized into finger-like protrusions of differentiated cells, called villi, and surrounding proliferative invaginations, called crypts of Lieberkühn.^[1,2]^ The intestinal epithelium undergoes fast and continuous cell renewal driven by the intestinal stem cells (ISCs) residing at the crypt bases. The progeny of the ISCs divide and differentiate while migrating up along the villi, forming distinct stem/proliferative and differentiated cell compartments.^[1,3]^ This compartmentalization is regulated by the interplay of gradients of biochemical factors, microbial metabolites, oxygen and extracellular matrix components formed along the crypt-villus axis.^[4–6]^ The biochemical factors essential for ISC niche maintenance, namely, Wingless-related integration site (Wnt), R-Spondin, and epidermal growth factor (EGF), are only active at the crypt compartments with a decreasing concentration from the crypt base towards the villi.^[5]^ These ISC niche factors arise from multiple sources; on one hand, from the epithelium itself, as Paneth cells found at the crypt bases are well known producers of Wnt3a and EGF factors.^[7]^ On the other hand, subepithelial myofibroblasts (ISEMFs) found below the crypts in the lamina propria have been shown to secrete canonical Wnt2b,^[8,9]^ R-Spondins^[10]^ and bone morphogenetic protein (BMP) antagonists (e.g. Noggin, Gremlin 1).^[11,12]^ Although it is known that the gradient of epithelial-autonomous Wnt3a is formed by plasma membrane dilution,^[13]^ it remains to be deciphered which is the nature of all remaining biomolecular gradients coming from the mesenchyme and how the epithelium integrates those signals.

Traditionally, it has been a challenge to grow intestinal epithelial cells in vitro. Therefore, mouse models have been fundamental in understanding the intestinal epithelial biology and the maintenance of the ISC niche in vivo. Lately, the development of organotypic, three-dimensional (3D) intestinal organoids^[14–16]^ has greatly improved the study of intestinal physiology and pathology in vitro.^[17,18]^ Intestinal organoids are derived from ISCs embedded in a basement membrane matrix (Matrigel), where they expand and self-organize, forming crypt-like and villus-like domains enclosing a lumen. They recapitulate the main cellular components and function of the native tissue. However, their closed geometry is a major limitation when using organoids to better understand the role of biochemical gradients in the tissue homeostasis or restoration. The surrounding 3D matrix prevents the spatiotemporal control of diffusive species present in the cell culture medium. In addition, classically cultured organoids are very heterogeneous in size and number of crypts and do not allow an easy access to the lumen. It is also acknowledged that the 3D native tissue architecture is fundamental to the formation and maintenance of biomolecular gradients, which subsequently, regulates cell compartmentalization in vivo.^[19]^ In this context, numerous attempts have been made to grow organoid-derived primary cells on microstructured natural hydrogels.^[20–22]^ However, so far it has been challenging to grow primary epithelial cells on synthetic materials which would provide a well-controlled microenvironment mimicking not only the native tissue 3D architecture but also including the spatiotemporal control of diffusive biochemical factors. This could be instrumental to understand how gradients of such factors regulate gut epithelial organization.

Microfabrication techniques provide a wide set of tools to better mimic the in vivo tissue architecture and function by allowing precise control over the cellular microenvironment.^[20,23–26]^ Few microengineered systems, reproducing intestinal and colonic architecture, and biomolecular gradients on collagen gels, have proven effective in inducing distinct stem/proliferative and differentiated cell compartmentalization in the monolayers obtained from primary cells.^[24,25,27]^ The colonic arrays obtained in this way were used as a screening platform for microbial metabolites and cytokines. However, the use of a multi-step labor-intensive strategy; natural hydrogels like collagen not being conducive to controlled modifications and being prone to degradation over time, even though reduced by crosslinking;^[20,24]^ and the possible unspecific interactions between the diffusing molecules and the collagen matrix hindering the gradient formation like in the case of rapidly absorbed hydrophobic drugs,^[28]^ pose certain limitations in downstream applications of such culture system. To be of practical use, the fabrication strategy should be easy to implement. Also, synthetic materials, such as poly(ethylene glycol) (PEG), have some advantages over the natural matrices, in allowing better control over biochemical and mechanical properties. In this way, their effect on cellular behavior can be systematically tested facilitating the development of an optimal, well-controlled microenvironment.

Here we employed an in-house-developed, simple and cost-effective photolithography-based technique to fabricate 3D villus-like poly(ethylene glycol) diacrylate (PEGDA)-based hydrogels^[29]^ with a controlled mesh size allowing the diffusion of ISC niche biomolecular factors. By developing a custom-made microfluidic device allocating the microstructured hydrogels, we imaged the gradients of a fluorescently labelled model protein formed along the vertical axis of the microstructures using a custom-made light sheet fluorescence microscope (LSFM).^[30]^ We then developed an in-silico model matching the experimental data and used the model to predict the gradients of ISC niche factors formed on the microstructures. We adapted these villus-like hydrogels to standard Transwell inserts that permitted access to both sides of the hydrogel, and subsequently, the simple formation of spatiotemporal gradients by using the lower compartment as the source and the upper compartment as the sink. We demonstrated the platform to be suitable for the growth, proliferation and differentiation of intestinal epithelial cells derived from organoids, thus showing for the first time the culture of these primary cells on microstructured PEG-based hydrogels. Through this platform, we also proved that the presence of biochemical gradients of ISC niche factors secreted by ISEMFs led to the in vivo-like compartmentalization of the proliferative and differentiated cells along the villus axis acting synergistically with the epithelial-derived gradients imposed by the architectural cues. Moreover, modifying the gradient profiles by changing the biochemical factor composition or the concentration range, as demonstrated by the simulations, affected the cellular distribution along the villus axis allowing us to relate specific gradient profiles to the cellular response over time. The physiologically representative and accurate platform that we present here, combining the 3D architecture and the gradients of ISC biochemical factors coming from ISEMFs, can be used to systematically test in vitro the effect of a wide range of biochemical factors that are essential for ISC maintenance and differentiation, especially those from intestinal stroma, on the intestinal cellular response and thus better understand the dynamics of the tissue.

## Results

The 3D villus-like microstructured hydrogel scaffolds were fabricated by lithography-based dynamic photopolymerization of PEGDA-AA prepolymer solutions with some modifications to the original protocol.^[29]^ First, the microstructures, and then, the hydrogel base in which they were sustained were formed on flexible porous PET membranes, ready to be assembled into Transwell inserts (**Figure 1 A**). The base thickness was 185 ± 24 μm and the villus-like microstructures were 350 ± 44 μm in height (Figure 1 A) resembling the anatomical dimensions of the small intestine villi.^[31–33]^ The hydrogels had a density of 16 villi mm^-2^, which is in good agreement with the density of the anatomical villi (20-40 villi mm^-2^).^[32,34]^ The hydrogel bulk elasticity measured for the samples was also consistent with the elastic properties of small intestinal tissue (**Figure S1** A).^[35]^

**Figure 1.**
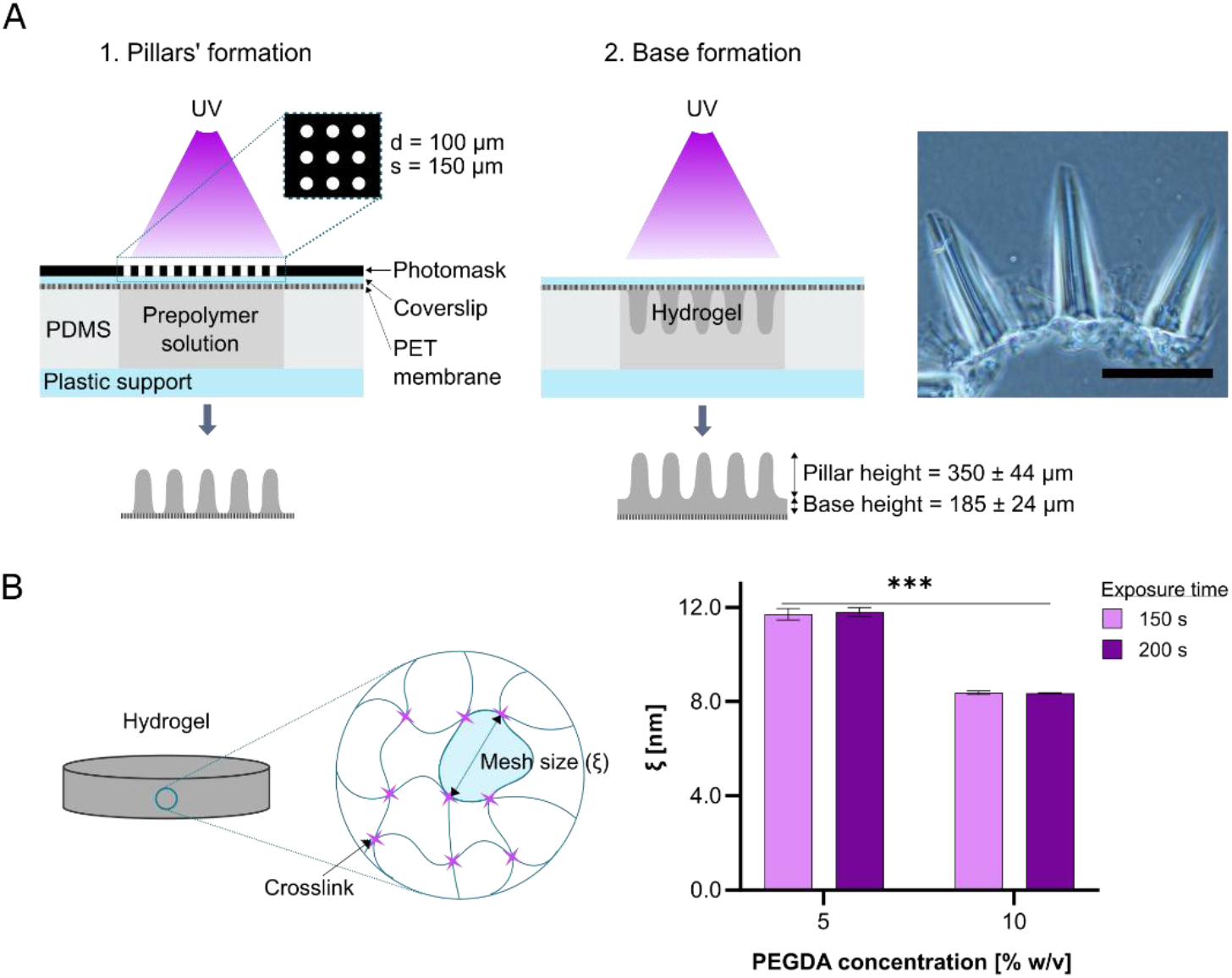
(A) Schematic drawing of the fabrication procedure of villus-like microstructured PEGDA-AA hydrogel scaffolds on porous PET membranes (left and middle). During the first exposure (left), the microstructures were formed using a photomask consisting of an array of circular transparent windows with a diameter (d) and a spacing (s) of 100 μm and 150 μm, respectively. In the second exposure (middle), the photomask was removed and the base with a 6.5 mm diameter (PDMS pool dimension) was formed. This step was needed to create a homogeneous hydrogel surface. Bright field microscope image of the cross-section of the villus-like microstructured hydrogel scaffolds (right). The height of the microstructures was measured to be 350 ± 44 μm. N=3, n=4, pillars=147. The height of the base was measured to be 185 ± 24 μm. N=4, n=5, pillars=110. Mean ± SD. Scale bar: 400 μm. (B) Schematic representation of the mesh size (*ξ*) (left). Graph showing estimated mesh size (*ξ*) for 5% and 10% w/v PEGDA-AA hydrogels fabricated with 150 s and 200 s of UV exposure (n=4) (right). P-value: ***p<0.001. Mean ± SD.

To have a better insight into the protein diffusivity within the network of PEGDA-AA hydrogels, the bulk mesh size (ξ) was estimated using Flory-Rehner equilibrium swelling theory^[36,37]^ as modified by Peppas and Merrill for hydrogels prepared in the presence of water^[38,39]^ (see Supporting Information). The estimated average mesh sizes changed from 11.7 ± 0.2 nm to 8.3 ± 0.2 nm, when the PEGDA concentration was increased from 5% w/v to 10% w/v, respectively (Figure 1 B), in accordance with the values reported in the literature.^[40,41]^ The tuning of the mesh size of the hydrogels by altering the PEGDA concentration is well established in the literature.^[42–44]^ A decrease in the average mesh size hinders the mobility of the polymer chains, limiting the diffusivity of the molecules through the hydrogel. For proper diffusion of the biomolecules, the mesh of the hydrogel network has to be equal to or greater than the diameter of the biomolecules. Taking into account that the hydrodynamic diameters of the main ISC niche biochemical factors (EGF, R-Spondin 1, Noggin, Wnt2b) range from 3.0 to 7.2 nm (see Table 1), the hydrogels with 5% - 10% w/v PEGDA concentration range are suitable for the diffusion of these proteins. On the other hand, there was no significant change in the mesh size for the two different UV exposure times studied, suggesting that maximum macromer conversion has already been achieved at 150 s of UV exposure.

**Table 1:** ISC niche biochemical factors and their corresponding model proteins with their molecular mass and the hydrodynamic diameters of the model proteins.

| ISC niche factor | MW [kDa] | Model protein | MW [kDa] | $d_h$ [nm] |
| --- | --- | --- | --- | --- |
| EGF | 6.0 | Insulin | 5.7 | 2.94<sup>[68]</sup> |
| R-Spondin 1 | 25.6 | CA<sup>a)</sup> | 29.0 | 4.70<sup>[68]</sup> |
| Noggin | 46.4 | BSA | 66.5 | 7.20<sup>[44]</sup> |
| Wnt2b | 38.0 | Ovalbumin | 43.0 | 5.96<sup>[68]</sup> |

<sup>a)</sup>CA: Carbonic anhydrase.

In order to be able to visualize gradients of ISC niche factors formed within villus-like microstructured hydrogels by microscopy techniques, a custom-made microfluidic device that allowed allocating the PEGDA-AA hydrogel was designed and fabricated. The SU-8 masters were fabricated by photolithography and replica molded in PDMS. The PDMS chip consisted of a microfluidic fluidic channel and a source chamber at the center, allocating the hydrogel sample right above the chamber (**Figure 2 A** left, **Figure S2** A). The hydrogel samples were fabricated on porous PET membranes and stuck to the microfluidic chip by double-sided pressure-sensitive adhesive (PSA) rings or by using uncured PDMS (Figure 2 A left, Figure S2 A). The fluorescently labelled protein solutions were delivered through the channel to the source chamber to generate the biomolecular gradients along the villus axis of the microstructured hydrogels. Fluorescently labeledgradients were visualized in-chip by LSFM.^[30]^ LSFM technique was selected as it permits fast imaging (minimal photobleaching) at large working distances with fine axial resolution, a trade-off difficult to achieve by traditional confocal microscopy.^[45,46]^ By decoupling the illumination and detection light paths, LSFM allows a flexible sample access and mounting, which was needed for the microfluidic device (Figure 2 A middle). The chip was designed in a way that the hydrogel scaffold would stick out allowing proper physical access required for imaging (Figure 2 A left, Figure S2 A). To fit the microfluidic chip in the microscope, a custom-made sample holder was designed and fabricated (Figure 2 A right). This had transparent walls permitting the entrance of the light-sheet to the sample and a pedestal with 25° inclination to have better access to the hydrogel and the source chamber allowing proper imaging (Figure 2 A right).

**Figure 2.**
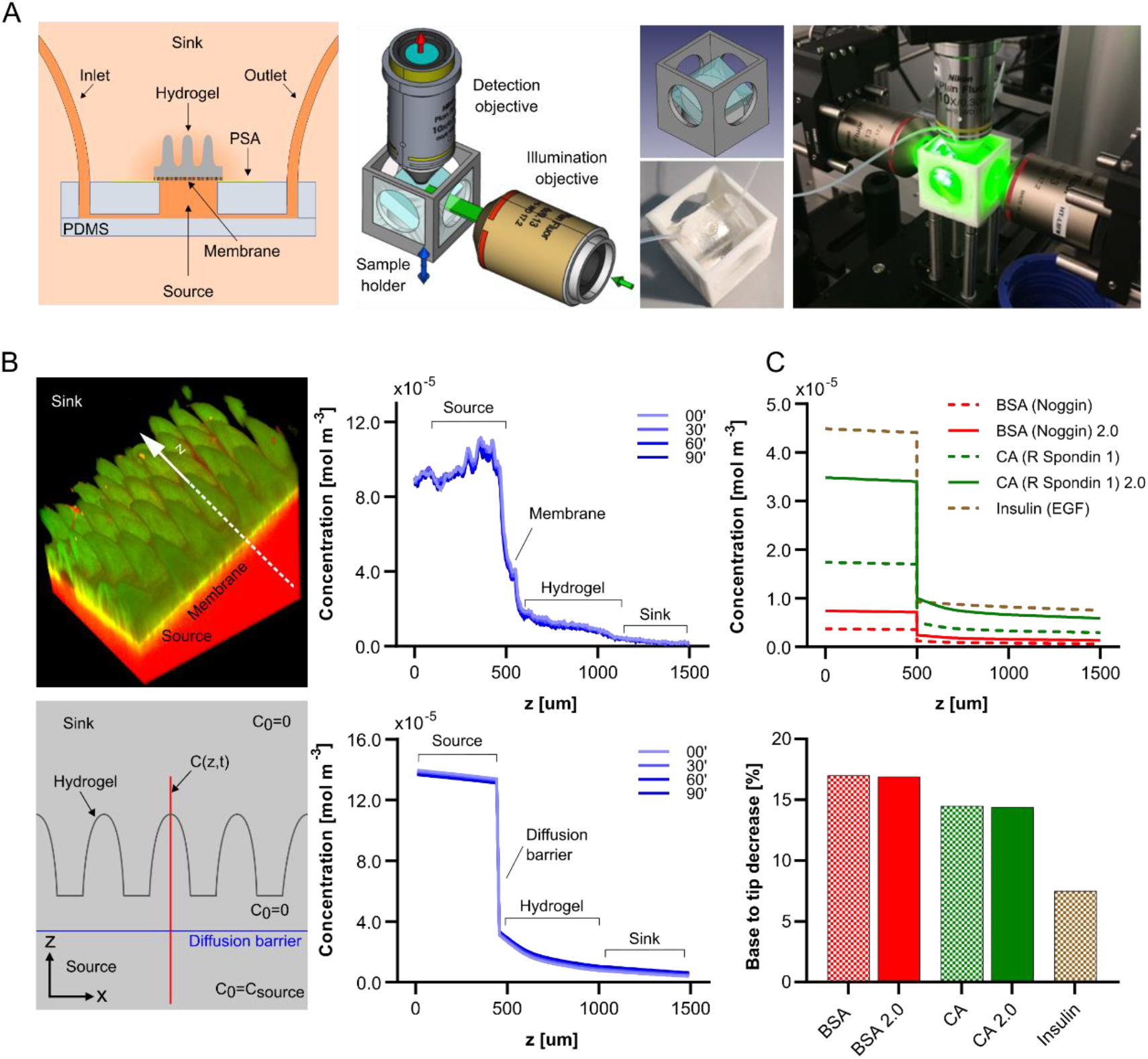
(A) Schematic drawing of the microfluidic chip allocating villus-like microstructured PEGDA-AA scaffold (left). The LSFM setup used, where the detection was performed in an up-right configuration (middle). Schematic of the imaging chamber designed, picture of the microfluidic chip glued to the imaging chamber and the final setup (right). (B) 3D reconstructed fluorescence image of Texas Red-labeled BSA (red) diffusion through villus-like microstructured scaffolds (green) acquired by LSFM spanning the source, the hydrogel and the sink at 3 hours after loading (top left). Graph showing the concentration profiles of Texas Red-labeled BSA taken along the white arrow shown in the left at different times points (top right). The beginning of the time-lapse was set as the zero time point. Samples from two independent experiments showed similar results. Scheme showing the microstructure geometry and the initial conditions used in the FEM simulations (bottom left). The initial concentrations were set to zero in the hydrogel and the sink, and to the corresponding experimental value in the source. The concentration profiles as a function of time along the red line spanning the source, the hydrogel and the sink were determined. The sink and source sizes are not to scale for visualization purposes. Graph showing the concentration profiles predicted by the FEM simulations of the diffusion of the model protein BSA (bottom right). (C) Graph showing the concentration profiles at the initial conditions predicted by the FEM simulations at 24 h for BSA (Noggin), BSA 2.0 (doubled source concentration), CA (R Spondin 1), CA 2.0 (doubled source concentration) and Insulin (EGF) taken along the red line shown in B bottom left (top). Graph showing the percent decrease in concentrations of the proteins from base to tip of the microstructures (bottom).

For the characterization of the biomolecular diffusion dynamics, Texas Red-labelled BSA was selected as the model protein. The diffusion through the microstructured hydrogel scaffolds was imaged by performing a time-lapse acquisition (Figure 2 B top left). The microfluidic chip allocating the hydrogel was loaded with Texas Red-labelled BSA and an overall 3D volume of 1.33 × 1.33 × 2.30 mm was imaged over 90 minutes. Besides Texas Red emission, we also acquired autofluorescence emission from the hydrogels, excited by a 488 nm laser, obtaining supplementary information about the hydrogel morphology and the membrane interface. The intensity profiles along the perpendicular axis of the hydrogels were taken and converted to concentration values by using a previously established calibration curve within the same setup (Figure 2 B top right). The concentration profiles showed that a gradient was generated within the hydrogels due to the diffusion of the proteins from the source to the sink. The gradient profiles were stable along the time of acquisition (Figure 2 B top right), indicating that 3 hours loading time were sufficient to reach a pseudo steady state leading to the formation of stable protein gradients within the hydrogels. On the other hand, the protein accumulation at the PET membrane-hydrogel interface was found to be significant and exact amount of protein retention was measured to be incorporated in the in silico model. During acquisition, the source and the sink were not perfused consistent with the entirety of the experiments performed in this work.

In silico modelling of biomolecular diffusion allows to predict the gradient profiles of proteins of interest that would have, otherwise, been tedious to determine experimentally. To that end, an in-silico model of diffusion was developed using finite element method (FEM). In agreement with the experiments performed, BSA was chosen as the model protein to develop the model, the microstructure geometry, finite volumes (no-flow condition) and the material accumulation at the membrane-hydrogel interface as measured in LSFM experiments were implemented (Figure 2 B bottom left). The concentration profiles computed along the perpendicular axis spanning the source, the hydrogel and the sink in the geometry designed matching the experimental setup also predicted stable gradients (a pseudo-steady state) within the timeframe studied, in perfect agreement with the experimental data (Figure 2 B bottom right, Figure S2 B). However, in a system with finite volumes, a pseudo-steady state (where change in flux is insignificant) can be maintained only within a specific time window. The system would equilibrate eventually unless the source and the sink are replenished. In fact, periodic replenishment of media allows the maintenance of the gradient profiles obtained at the periodicity.^[47,48]^ To demonstrate this point, the periodic medium replenishment performed every 24 h (convenient periodicity for manipulation purposes) was implemented in the model. The simulations showed that, indeed, this strategy allowed to establish stable gradients (**Figure S3**), and therefore, it was implemented in the cell culture experiments. Once the in-silico model was validated, it was used to predict the initial conditions of the concentration gradients of ISC niche biochemical factors Noggin, R-Spondin 1, and EGF formed along the villus axis of the hydrogels in the cell culture setup (Transwell system), using data from the model proteins BSA, CA, and Insulin, respectively (Figure 2 C top). Simulations revealed that when the source concentrations were doubled (BSA 2.0 and CA 2.0), the concentrations in the entire geometry were also doubled. On the other hand, the percent decrease from base to tip were found to be to be 17.0%, 16.9%, 14.5%, 14.4%, and 7.5% for BSA, BSA 2.0, CA, CA 2.0 and Insulin, respectively (Figure 2 C bottom), showing that relative change is unaffected by changes in the source concentration. Therefore, our system enables to regulate the absolute concentrations that the cells can sense, while keeping the relative drop constant.

After characterizing the gradients of ISC niche factors formed on PEGDA-AA villus-like hydrogels by LSFM and in-silico modelling, we assessed whether we could grow complete organoid-derived epithelium monolayers on the microstructured hydrogels bearing the biochemical gradients. For that, the PEGDA-AA villus-like hydrogels were fabricated on PET membranes, mounted on Transwell inserts and functionalized using collagen type I (**Figure 3 A**). Then, the proteins were delivered in the Transwell insert asymmetrically to create the spatial gradients. For this first step, we employed the cocktail of factors we use to culture organoids (ENRCV) supplemented to conditioned medium from intestinal subepithelial myofibroblasts (ISEMF_CM) and we corrected the factor concentrations for the protein accumulation taking place in the membrane as predicted by the in-silico simulations. 24 hours after the formation of the gradients, we seeded organoid derived single cells from mouse isolated intestinal crypts. With the aim of optimizing the formation of the epithelial monolayers, we first tested the scaffolds fabricated with microstructures separated by either 100 µm or 150 µm and found that 150 µm spacing provided a sufficient base area for the epithelial monolayer to be formed (**Figure S4** A). Also, among the AA content tested in the hydrogel base formulation, it was found that 0.3% w/v AA allowed the formation of monolayers with characteristic epithelial morphology at the base region (Figure S4 B). Finally, we found that, for most crypt isolations, the minimum cell seeding density required to obtain a full epithelium on the scaffold at day 3 was 5×10^5^ cells/sample (Figure S4 C).

**Figure 3.**
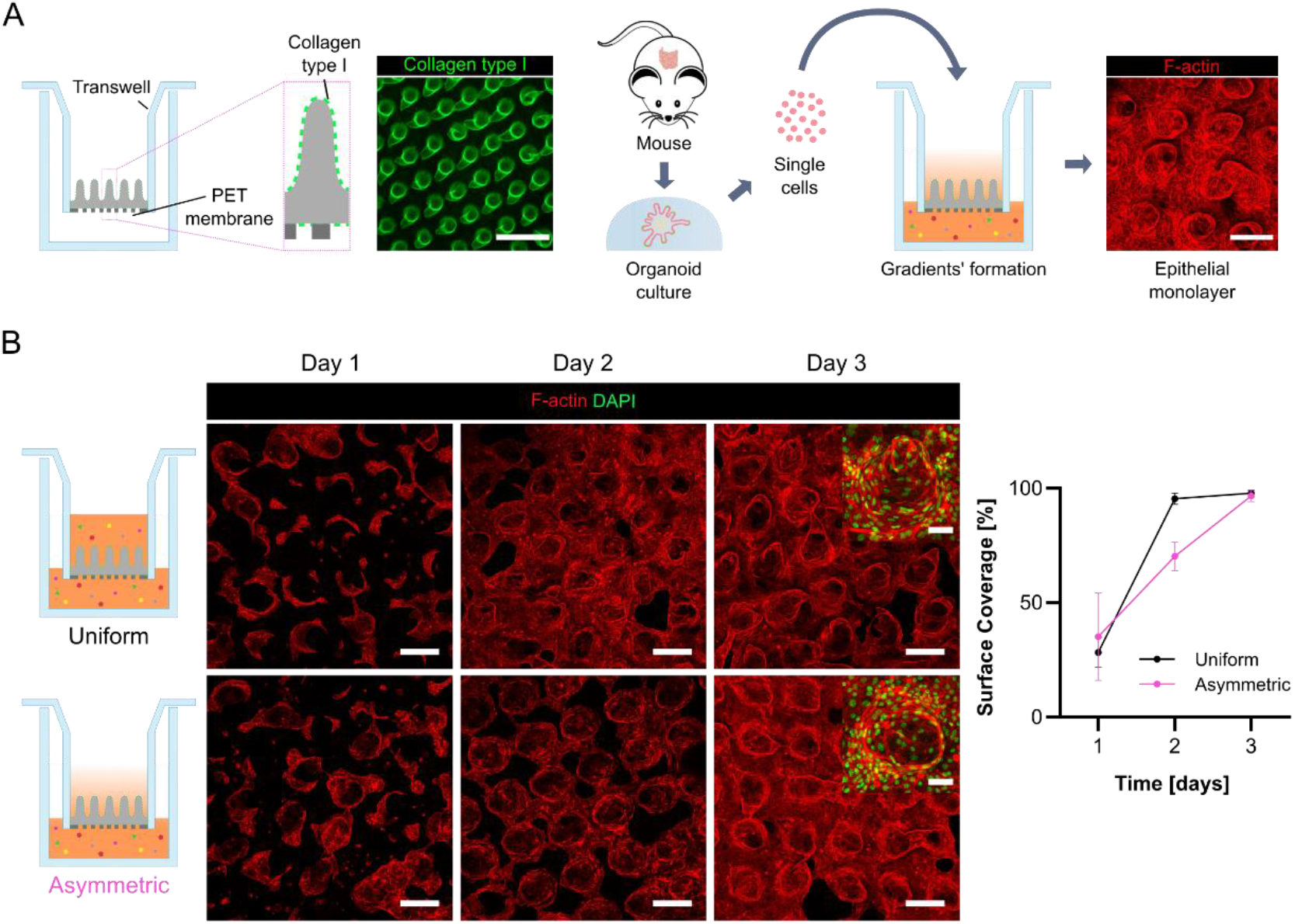
(A) Schematic drawing showing a scaffold mounted onto a Transwell insert and functionalized with collagen type I, an image of a functionalized scaffold, a schematic drawing showing the organoid-derived single cell seeding on collagen functionalized scaffolds with the gradients of ISC niche proteins already generated and an image of the epithelial monolayer formed (from left to right). (B) Schematic drawings of Uniform and Asymmetric conditions. Representative confocal maximum intensity projection images of samples at day 1, 2 and 3 of culture immunostained for F-actin. Scale bars: 200 µm. Zoom-in insets at day 3 immunostained for F-actin and DAPI. Scale bars: 50 µm. Graph plotting the percentage of surface covered by epithelial cells with respect to the total surface as a function of the cell culture time in the absence (Uniform) or presence (Asymmetric) of ISC biochemical gradients. Asymmetric: Day 1 N = 3, n = 3, Day 2 N = 2, n = 2, Day 3 N = 4, n = 4. Uniform: Day 1 N = 3, n = 3, Day 2 N = 2, n = 2, Day 3 N = 3, n = 3. Mean ± SD.

Having established the optimal conditions for the growth of complete epithelial monolayers, we wondered if the process of monolayer formation was dependent on the presence or absence of ISC niche biomolecular gradients. For this purpose, we seeded organoid-derived single cells on top of scaffolds bearing gradients and on scaffolds with no gradients, that is to say, with uniform concentration of the factors. Every 24 hours we renewed the media of both apical and basolateral compartments to maintain the gradients stable (**Figure S3**). One day after cell seeding, cells were starting to form a monolayer, covering approximately 30% of the surface under both, asymmetric and uniform conditions (Figure 3 B). 48 hours after seeding, the monolayer extended to 70% of the surface in the asymmetric condition while in uniform condition, the surface was almost fully covered. At 72 hours, the cultures reached nearly 100% surface coverage in both conditions (Figure 3 B). We hypothesize that this faster progression of single cells into full monolayers under uniform condition compared to asymmetric might be due to the greater availability of stem cell niche factors in this condition as factors are not only delivered basolaterally but also apically. Still, it appears that organoid-derived cells were able to form complete monolayers on the surface of the hydrogels regardless of the presence or absence of such gradients. This implies that the type of ISC niche factor gradients formed in our system were not necessary for epithelial cells to be able to form complete epithelial monolayers; provided that the minimum concentration of factors required for cell growth were supplied. Still, it remained to be deciphered whether gradients of ISC factors determine cell compartmentalization in vitro.

For that aim, we analyzed the location of proliferative (Ki67^+^), stem (GFP^+^), and differentiated (CK20^+^) cells including Paneth cells (Lyz^+^) and enterocytes (FABP1^+^) in the presence or absence of gradients of ISC niche factors when there was a complete epithelium (i.e. at day 3). First, we analyzed the compartmentalization of proliferative Ki67^+^ cells along the villus axis. Under gradients of stem cell niche factors, the frequency of proliferative cells positioned at the base was significantly higher (**Figure 4 A**). We observed that almost 50% of proliferative were located at the base, while only 11% were at the tip of the pillars. Instead, when we administered the factors uniformly without creating gradients, less than 30% of the Ki67^+^ cells were located at the base, and above 15% were located at the tip and the difference was not statistically significant (Figure 4 A). The quantification of the GFP signal along the villus axis revealed that GFP^+^ stem cells were also preferentially located at the base in the asymmetric condition, while they were rather uniformly dispersed among the whole axis of the pillar in the uniform condition at day 3 (Figure 4 B, **Figure S5** A). Most of the Ki67^+^ cells colocalized with GFP^+^ stem cells in both conditions at day 1, while at day 3 68.2 ± 5.0 % and 93.7 ± 2.0 % of Ki67^+^ cells were also GFP^+^ for asymmetric and uniform conditions, respectively (Figure S5 B). This shows that the stem cell domain is smaller in the asymmetric condition, presumably due to the cells receiving less ISC niche factors. On the other hand, lysozyme (Lyz)^+^ Paneth cells were also preferentially located at the base for both conditions (Figure S5 C) and they were colocalizing with GFP^+^ stem cells. Altogether, these findings suggest that the presence of gradients guides the formation of crypt-like compartments containing proliferative, stem and Paneth cells to localize at the base of the pillars. Moreover, to assess the evolution of this in vivo-like compartmentalization of proliferative cells in culture, we quantified their relative change of frequency at the base and at the tips from day 1 to day 3. We observed that under asymmetric administration of the factors, the frequency of Ki67^+^ cells at the base increased significantly with time, while decreasing over the pillars (Figure S5 B), suggesting that the concentrations of factors sensed by the cells located at the pillars were not enough for them to maintain the proliferative state. Although a similar tendency was observed under the uniform administration of the factors, the difference was much fainter and not statistically significant (Figure S5 B). Then, we wanted to assess how the positioning of the differentiated population in complete epithelium state was impacted by the gradients of stem cell niche factors. For that, we analyzed the mean intensity of CK20, a marker for differentiated cells with a graded as opposed to binary expression.^[49]^ In the absence of the gradients, the normalized mean intensity of CK20 was similar all along the pillars. In contrast, in the presence of gradients, there was a steep and statistically difference of such intensity, being lower at the base and higher at the tip of the pillars (Figure 4 C, Figure S5 A). Similarly, the amount of fatty acid binding protein 1 positive (FABP1^+^) enterocytes concentrated at the tip of the pillars was significantly higher in the presence of gradients, as opposed to the uniform condition (Figure 4 D). On the other hand, we detected few Periodic acid-Schiff base (PAS)^+^ goblet cells dispersed along the pillars for both conditions (Figure S5 D). Altogether, these findings suggest that the presence of gradients favors the in vivo cell populations and their more physiological disposition.

**Figure 4.**
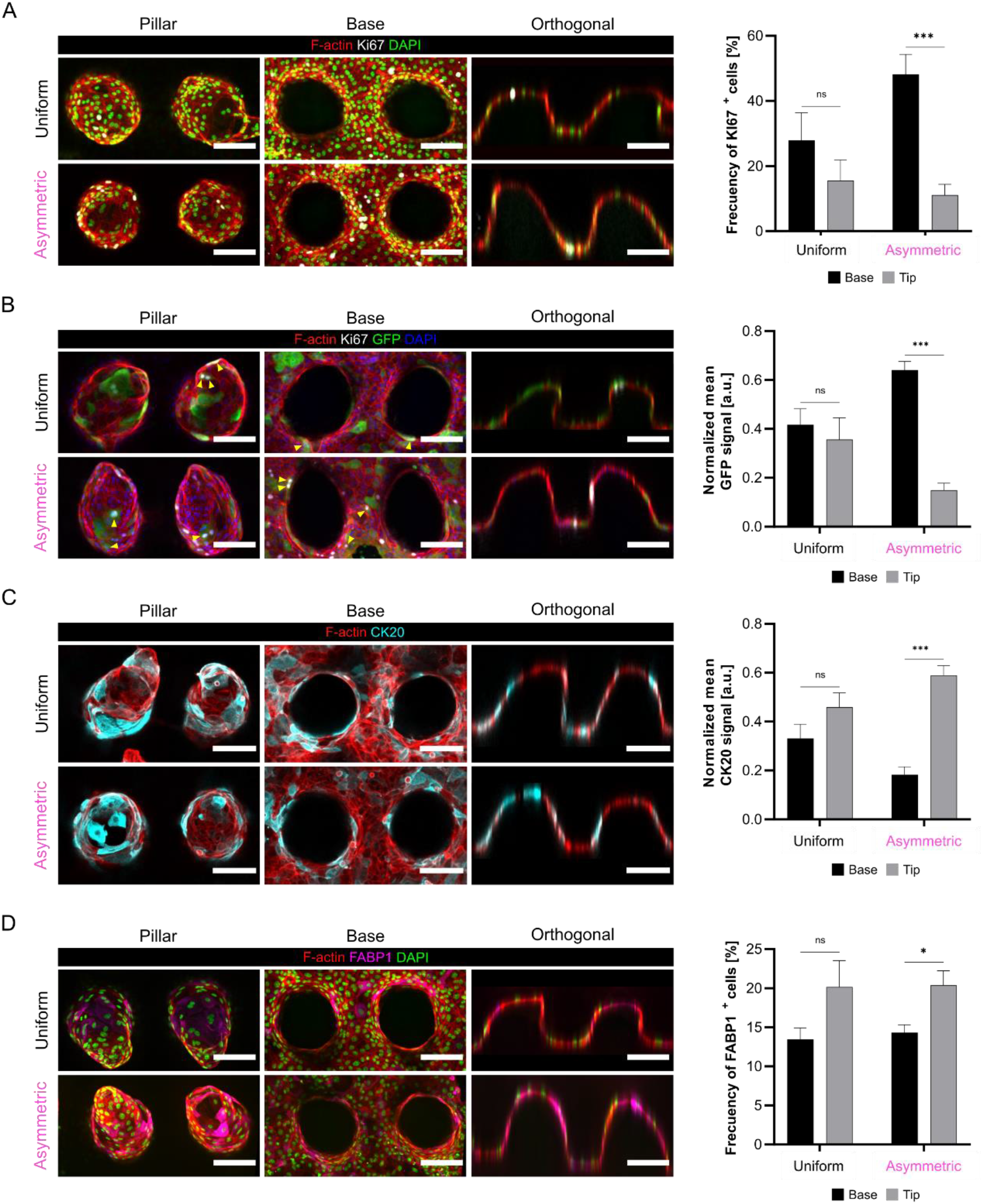
Maximum intensity projections of the sections corresponding to the pillar (left) and to the base (middle) and orthogonal section of the whole villus axis (right). Representative images of Uniform (top) and Asymmetric (bottom) samples on day 3 of culture. (A) Samples immunostained for F-actin, Ki67, GFP (stem cells) and DAPI. Frequency of Ki67^+^ cells at day 3 of culture at the base and the tip of the pillars with respect to the total Ki67^+^ cells under ISC-niche biomolecules gradients (Asymmetric) or no gradients (Uniform) (left). Uniform: N = 2, pillars = 19. Asymmetric: N = 4, pillars = 41. P-value: *** < 0.001. Mean ± SEM. (B) Samples immunostained for F-actin, Ki67 (proliferative cells) and DAPI and F-actin Ki67, GFP (stem cells) and DAPI. Normalized mean GFP signal at the base and the tip of the pillars in Asymmetric or Uniform conditions at day 3 of culture (right). Uniform: N = 1, pillars = 12. Asymmetric: N = 1, pillars = 12. P-value: *** < 0.001. Mean ± SEM. (C) Samples immunostained for F-actin and CK20 (differentiated cells). Normalized mean CK20 signal at the base and the tip of the pillars in Asymmetric or Uniform conditions at day 3 of culture (right). Uniform: N = 2, pillars = 19. Asymmetric: N = 4, pillars = 41. P-value: *** < 0.001. Mean ± SEM. (D) Samples immunostained for F-actin, FABP1 (enterocytes) and DAPI. Frequency of FABP1^+^ cells at day 3 of culture at the base and the tip of the pillars with respect to the total FABP1^+^ cells under ISC-niche biomolecules gradients (Asymmetric) or no gradients (Uniform) (left). Uniform: N = 1, pillars = 13. Asymmetric: N = 1, pillars = 11. Two-tailed t-test. Mean ± SEM. P-value: * < 0.05.

After observing that the gradients of stem cell niche factors formed in our in vitro platform modulate the proliferative and differentiated cell positioning, we made use of our 3D in vitro system to modify the gradients applied and to read out the effect on the cellular response. For this purpose, we devised two new conditions: in the first one, we doubled the concentrations of Noggin and R-Spondin 1 (Asymmetric 2.0); and in the second one, we tested a chemically defined medium by plainly substituting ISEMF_CM in Asymmetric 2.0 for Wnt2b, the most relevant proliferation-promoting molecule coming from the mesenchyme^[9,50]^ (Asymmetric Wnt2b) (Table 1, Table S1). The in-silico model we developed predicted that doubling the source concentration of R-spondin 1 and Noggin would result in doubled concentrations for a given point along the villus axis, while the gradient slopes would stay unchanged (**Figure 5 A**). Cells receiving doubled amounts of these ISC niche factors account for the increased expansion of the epithelial monolayers we observed at day 1 (Figure 5 B). The surface coverage (mean ± SEM) was 84.2 ± 5.6% and 83.7 ± 6% for Asymmetric 2.0 and Asymmetric Wnt2b conditions, respectively, compared to 50.9 ± 17.6% for the Asymmetric condition. At day 3, epithelial cells occupied almost the entire scaffold surface for all the conditions (96.2 ± 0.9% for Asymmetric, 97.2 ± 1.1% for Asymmetric 2.0 and 93.1 ± 3.6% for Asymmetric Wnt2b), therefore constituting a full epithelial monolayer. The faster expansion of the epithelial monolayers under both Asymmetric 2.0 and Asymmetric Wnt2b conditions can be explained by the increased amounts of proliferative cells. At day 1, the percentage of Ki67^+^ proliferative cells was significantly higher in Asymmetric Wnt2b compared to the other two conditions (Figure 5 C top). On the other hand, the frequency of Ki67^+^ cells was significantly reduced at day 3 for the same condition, suggesting that the type of Wnt2b gradient present in our system leads to an initial fast expansion and a later exhaustion of the proliferative domain, altering the growth rhythm of the epithelial monolayer.

**Figure 5.**
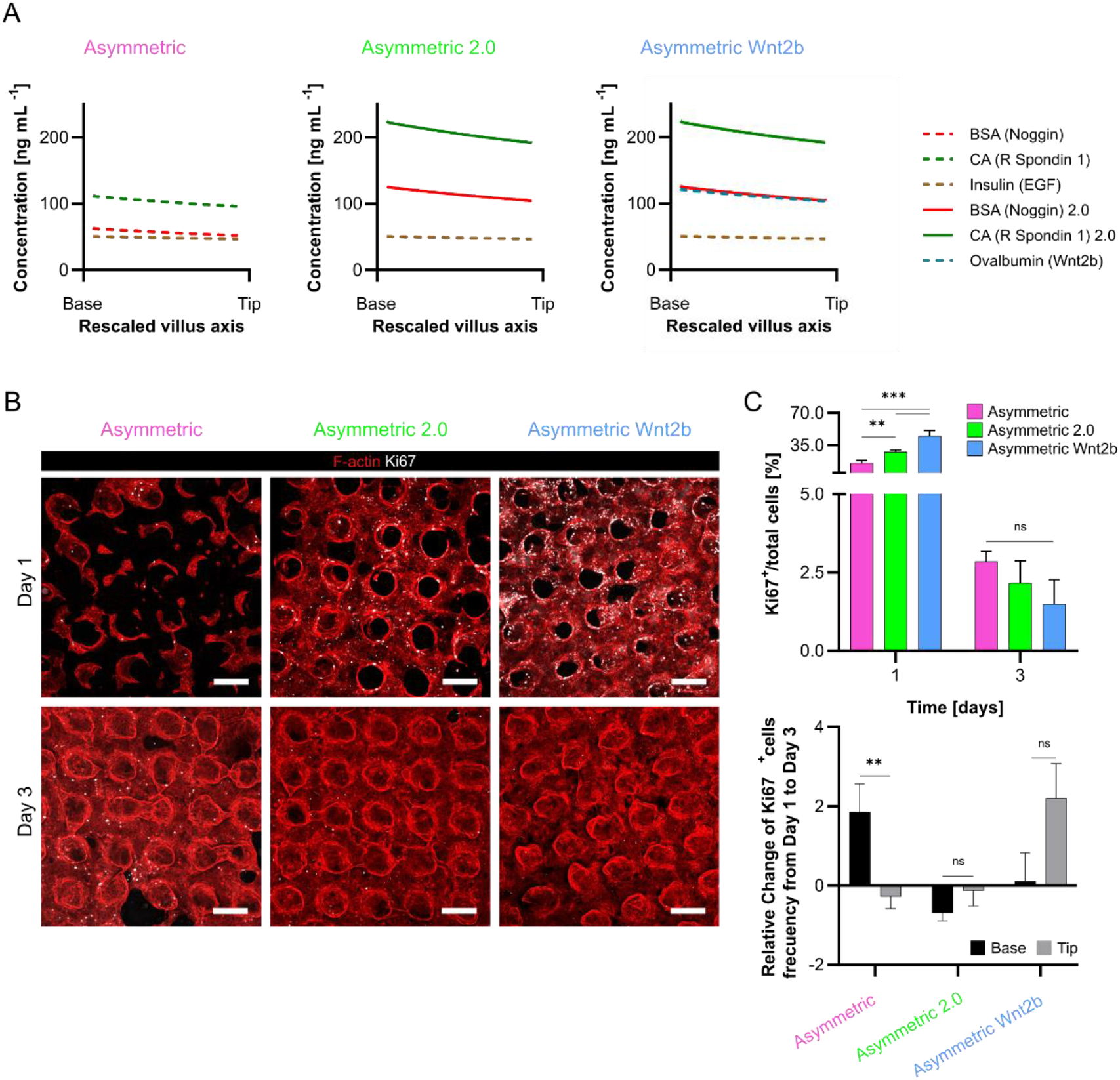
(A) Graphs showing the change in concentrations as a function of position along the villus axis predicted by the FEM simulations at 24 h for BSA (Noggin), BSA 2.0 (doubled source concentration), CA (R Spondin 1), CA 2.0 (doubled source concentration), Insulin (EGF) and Ovalbumin (Wnt2b) present in the corresponding Asymmetric (left), Asymmetric 2.0 (middle), Asymmetric Wnt2b (right) culture conditions. (B) Representative confocal maximum intensity projections images of Asymmetric, Asymmetric 2.0 and Asymmetric Wnt2b samples at days 1 and 3 of cell culture immunostained for F-actin and Ki67 (left). Scale bars: 200 µm. (C) Quantification of the percentage of Ki67^+^ (proliferative) cells with respect to the total number of cells for each culture condition (top). Asymmetric: Day 1, N = 3, pillars = 19, Day 3, N = 4, pillars = 40. Asymmetric 2.0: Day 1, N = 2, pillars = 13, Day 3, N = 2, pillars = 15. Asymmetric Wnt2b: Day 1, N = 2, pillars = 10, Day 3, N = 2, pillars = 15. P-value: ** < 0.01, *** < 0.001. Mean ± SEM. Quantification of the relative change of Ki67^+^ cells from Day 1 to Day 3 at the base and pillar sections for each culture condition (bottom). Asymmetric: N = 3, pillars = 19. Asymmetric 2.0: N = 2, pillars = 13. Asymmetric Wnt2b: N = 2, pillars = 10. Two-tailed t-test. Mean ± SD. P-value: ** < 0.01.

When we looked at the distribution of the proliferative cells under different gradients, we observed that the frequency of the proliferative cells significantly increased at the base compared to the tip from day 1 to day 3 under the Asymmetric condition, while it stayed unchanged at the base and the tip for Asymmetric 2.0 (Figure 5 C bottom). This finding suggests that, in Asymmetric 2.0 condition, the whole range of concentrations of the ISC factors from the base to the tip of the microstructures is high enough to maintain all the cells at proliferative state. However, in the Asymmetric condition, only the concentrations present at the base allow the cells to stay proliferative. Under Asymmetric Wnt2b condition, the frequency of the proliferative cells found at the base increased from day 1 to day 3 compared to Asymmetric 2.0, presumably due to replacement of ISEMF_CM for Wnt2b (Figure 5 C bottom). On the other hand, the distribution of proliferative cells was not significantly different between the base and the tip, in line with our results for Asymmetric 2.0 condition. Altogether, these findings demonstrate that different ISC niche biochemical gradients lead to changes in cell type composition and distribution in our villus-like 3D intestinal epithelial model, proving it useful for the study of such gradients on the cellular response.

## Discussion

Biomolecular gradients present along the crypt-villus axis of the intestinal epithelium play a key role in the tissue organization and homeostasis by driving the maintenance of the intestinal stem cells (ISCs) at the crypts and their ordered differentiation along the villus axis.^[51]^ The characteristic 3D architecture of the intestinal epithelium is of fundamental importance in vivo as it provides a physical template for the gradients to be formed and maintained.^[19]^ Still, intestinal organoids are able to self-organize into crypt and villus-like regions,^[14]^ as well as organoid-derived 2D monolayer models,^[52,53]^ thus, underlining the capacity of epithelial-derived gradients, mainly orchestrated by Paneth cells,^[7,13,54]^ to organize the tissue. Therefore, the decoupling of the intestinal geometry from the biochemical gradients present within is not straightforward and has not been achieved so far. Intestinal organoids that, otherwise, show stochastic growth with crypt buddings variable in number, size, and orientation, can be patterned in a predefined manner using microengineered systems as they provide cells with architectural cues,^[20,21,24]^ which, at their turn, provide a template for the biochemical cues.^[24,25]^ Since biomolecular gradients are spatially patterned through the crypt-villus axis, in vitro models need to provide a crosstalk between the architectural cues and the biomolecular gradients to induce tissue patterning and thus be physiologically relevant. Not only that, but also the gradients formed by biochemical factors such as Noggin and Gremlin,^[12]^ R-Spondins^[55]^ and Wnt2b^[9]^ secreted by intestinal subepithelial myofibroblasts (ISEMFs) located below the crypts are needed for ISC maintenance and differentiation and thus, need to be considered when engineering more physiologically relevant intestinal epithelium platforms.

Here, we used an in-house developed photolithography-based approach to produce 3D villus-like hydrogels^[29]^ to provide cells with topographical cues with well-controlled biochemical gradients, as characterized by the state-of-the-art light sheet fluorescence microscopy and the in-silico modelling. We employed a synthetic and non-degradable PEG-based hydrogel to fabricate the 3D villus-like scaffold since its inert nature minimizes unspecific interactions with diffusing molecules.^[28]^ In the studies published so far, both PEG-based synthetic^[56]^ and natural hydrogels like collagen,^[20,24]^ Matrigel^[21,22]^ or a mixture of the two^[21]^ were shown to support ISC growth; yet so far only the natural ones have been chosen as scaffold materials.^[20,21,24]^ However, collagen and Matrigel are susceptible to protease remodeling thus being liable to degradation over time, even though it can be slowed down by crosslinking.^[20,22]^ Therefore, the use of non-degradable and inert PEG-based hydrogels posed as ideal for systematic screening of diffusible factors in 3D in vitro models. On the other hand, one other favorable property of synthetic hydrogels like PEG is the ease of tuning their mechanical properties. The PEG-based hydrogels used in this study showed bulk elasticity ranging from 2.5 ± 1.1 kPa (PEGDA 5% w/v) to 10.9 ± 1.7 kPa (PEGDA 10% w/v), in accordance with the elasticity values reported in the literature for optimal ISC growth and differentiation (neutralized collagen = 0.1-1 kPa,^[24]^ crosslinked collagen = 9.46 kPa,^[24]^ crosslinked Matrigel = 418.8 Pa,^[22]^ polyacrylamide = 0.1-15 kPa^[22,57]^). And finally, the simple and moldless fabrication procedure that we adopted, as opposed to the replica molding approach that requires multiple steps and intermediate molds,^[20,24]^ enables to routinely incorporate 3D architectural cues in cell culture systems like Transwell inserts, which is at the same time an ideal tool to generate biochemical gradients as it provides two separate basolateral and apical compartments. Leveraging this, we established biochemical gradients of the ISC niche factors generated by ISEMFs and showed that these gradients were acting synergistically with the epithelial-derived gradients imposed by the 3D architecture in inducing an in vivo-like compartmentalization of proliferative and differentiated cells.

The use of native tissue-like microstructured scaffolds allows deciphering how architecture alone affects cellular compartmentalization in the epithelium formed. To our knowledge, it does so by shaping the gradients of the epithelium-derived signaling^[19]^ and by providing geometric cues that are necessary for the tissue mechanics e.g. confinement needed to impose apical constriction of the stem cell domain.^[57]^ For instance, scaffolds with crypt-like structures (diameter = 50-100 µm, height = 130-170 µm)^[21,22,24]^ were shown to enable the patterning of the crypt-like domains even in the absence of exogenously controlled biomolecular gradients,^[21,22]^ presumably due to the reasons given above. In our work, however, we show that even in the absence of the crypt-like structures but in the presence of gradients, stem and proliferative cells localize predominantly to the base region, closer to the source of the factors secreted by ISEMFs, whereas differentiated cells, mainly enterocytes, localize at the tips. This shows that architectural cues and gradients of factors coming from ISEMFs act synergistically. Encapsulating fibroblasts into the microstructured scaffolds is another strategy to study this interplay. The presence of fibroblasts was shown to improve the epithelial organization and function,^[20]^ through both cell-cell interactions and paracrine signaling,^[50,58,59]^ possibly acting as gradient sources. These systems, however, lack the control over the concentrations of each individual factor and the possibility of correlating the dose or the gradient profiles of relevant factors to the cellular response observed. A reductionist system, combining 3D architecture with well-controlled gradients, on the other hand, would allow to systematically screen factors and understand how, for instance, those coming from the stroma act on epithelial dynamics. To that end, using our platform, we showed that when we modified the gradient profiles by increasing concentrations or by adding new factors like Wnt2b, we no longer observed cell compartmentalization. We hypothesize that this loss of compartmentalization might be due to the higher concentrations that cells were receiving, as the gradients of ISC niche factors, which are only active in the crypt compartment in vivo,^[51]^ were spanning the entire vertical axis of the scaffolds. Therefore, using our system, one can determine which ISC niche components and their optimal source concentrations are needed to obtain in vivo-like tissue compartmentalization. Considering that the multiscale coupling of geometric and biochemical cues is a concerted effort in the field of synthetic morphogenesis/engineering organoids^[60,61]^ and there remain open questions, we believe that the culture platform that we present here, with physiologically representative and well-controlled microenvironment (3D architecture and gradients of ISC biochemical factors coming from the intestinal stroma), will be useful to systematically test the effect of a wide range of ISC niche factors in the context of understanding intestinal tissue dynamics. For instance, in a pathological context of injured intestinal mucosa, present in inflammatory bowel disease (IBD), it has been recently suggested that factors produced by the intestinal stroma are essential for the successful restoration of the damaged epithelium.[62–64] However, there is currently no knowledge of the individual contributions of each of the factors secreted by the fibroblasts. In this context, our platform could be used to pinpoint which are the essential ISC niche factors that drive the biological response of wound healing and restoration of homeostatic cell compartmentalization. We believe that our platform in this sense could be a useful tool for new preclinical studies.

## 4. Experimental Section

### Fabrication of villus-like microstructured hydrogels

The villus-like microstructured hydrogels were fabricated by photolithography, as previously described.^[29]^ Briefly, a prepolymer solution containing PEGDA (6.5% (w/v), 6 kDa of MW, Sigma-Aldrich), acrylic acid (AA) (0.3% (w/v), Sigma-Aldrich), and Irgacure D-2959 photoinitiator (1% (w/v), Sigma-Aldrich) in phosphate-buffered saline (PBS) was flown into a chip fabricated with a 1 mm thick polydimethylsiloxane (PDMS) (Sylgard 184, Dow Corning) stencil containing an array of pools of 6.5 mm diameter. Tracketch polyethylene terephthalate (PET) membranes of 5 μm pore size (Sabeu GmbH & Co.) were used as substrates for the hydrogels. The microstructured PEGDA-AA hydrogels were fabricated by exposing the pre-polymer solution to UV light under patterned photomasks with transparent windows of 100 μm in diameter, edge to edge spacing of 150 μm and a density of 16 windows mm^-2^ (designed by Inkscape software and printed on acetate films (CAD/Art Services)). The photolithography was performed in an MJBA mask aligner (SUSS MicroTec) using a power density of ≈ 25 mW cm^-2^, the exact power density was measured at every fabrication using a 365 nm probe (UV1000, SUSS MicroTec). The prepolymer solution was exposed to ≈ 220 s, the exact time being adjusted based on the power density of the lamp at every fabrication to obtain a total energy of 5.4 J cm^-2^ to form the villus-like micropillars. A second exposure of ≈ 25 s, the exact time being adjusted based on the power density of the lamp at every fabrication to obtain a total energy of 0.6 J cm^-2^, was performed to form a hydrogel base holding the microstructures together (Figure 1 A). After UV exposure, unreacted polymer and photoinitiator were washed out with PBS and the hydrogels were kept submerged at 4°C for at least 3 days to reach equilibrium swelling. The hydrogels dimensions were measured using a bright field microscope. First, rows of the hydrogel were cut and knocked down so as to have them flat to the surface. Next, pictures of different rows within different hydrogels were taken and the parameters height of the pillars, tip diameter, base diameter and thickness of the base were measured using Fiji software.

Samples fabricated onto PET membranes were then assembled on modified Transwell inserts or microfluidic chips using double-sided pressure-sensitive adhesive (PSA) rings (Adhesive Research) as detailed previously.^[29]^ After swelling, and to provide the scaffolds with cell-adhesion motifs, the PEGDA-AA hydrogels were functionalized with collagen type I (10% w/v or 0.4 mg mL^-1^, Sigma-Aldrich) via N – (3-Dimethylaminopropyl)-N ′ - ethylcarbodiimide (EDC)/N-Hydroxysuccinimide (NHS) (Sigma-Aldrich) mediated coupling. Right after functionalization, the samples were used for cell culture experiments.

### Characterization of the mesh size of PEGDA-AA hydrogels

The mesh size of PEGDA-AA hydrogels was estimated using Flory-Rehner equilibrium swelling theory as modified by Peppas and Merrill for hydrogels diluted in aqueous media.^[38,39]^ This was an essential parameter to assess the diffusivity of biomolecules through the hydrogels. Hydrogels were produced from pre-polymer solutions containing PEGDA (5 – 10 % (w/v)), I2959 (1% (w/v)) and AA (0.06% (w/v)) in PBS and were casted as discs, 10 mm in diameter and ≈ 1 mm in height. Note that changes in AA concentration from 0.06% to 0.6% w/v don’t have any significant effects on the swelling behavior of the hydrogels.^[29]^ Hydrogels were fabricated using a power density of 25 mW cm^-2^ and 150 s and 200 s of UV exposure. The weight of the PEGDA-AA hydrogels was then measured at three different time points, i.e. after fabrication (also called after curing, m_c_), at equilibrium swelling (swollen in PBS at 4°C, up to 7 days, m_s_), and at dry state (dried till constant weight, m_d_). Once these values were obtained, the mesh size was calculated using previously established equations.^[65]^ For the detailed explanation of the calculations and the equations used, see Supporting Information.

### Fabrication of a microfluidic chip for biomolecule gradient generation and visualization

A PDMS chip for the generation and visualization of biomolecular gradients through the hydrogel network was designed and fabricated. It contained a microfluidic channel and a chamber allocating individual hydrogels (Figure 2A, left). For the fabrication of the microfluidic channel, a mold was prepared using SU-8 resist on a silicon wafer (Si(111), 1-side polished, n-type) (MicroChemicals) and photolithography. The wafer was sequentially cleaned with acetone and isopropanol, and oxygen plasma cleaning (PDC-002, Harrick Scientific) for 20 min at low RF (6.8 W). After drying (hot plate at 95°C) to remove the excess of humidity, a 100 μm thick layer of SU-8 2100 negative photoresist (Microchem) was spun-coated (WS-650MZ-23NPP/LITE, Laurell Technologies) on the wafer. The film was baked for 5 min at 65°C and 20 min at 95°C to evaporate the solvent and it was exposed to UV light (25 mW cm^-2^) through a photomask with the channel design (in a rectangular transparent window, 200 μm in width and 20 mm in length). After a post-bake (5 min at 65°C and 10 min at 95°C), the SU-8 film was developed (Microchem developer), rinsed with isopropanol, and dried under nitrogen. Right after fabrication, the height of the channel was measured by profilometer (DEKTAK 6M, Veeco Instruments) as 104.3 ± 0.1 μm. To avoid sticking when producing PDMS replicas, SU-8 molds were silanized with (Tridecafluoro-1,1,2,2-tetrahydrooctyl)trichlorosilane (ABCR GmbH & Co. KG), in vapor phase. Then, PDMS replicas containing the microfluidic channel were punched with a 0.75 mm punch (Harris Uni-Core, GE Healthcare Life Sciences) at each end of the channel, and with a 4 mm punch at the center to form a chamber below the hydrogel (Figure S2 A). Afterwards, they were plasma bonded to a 1 mm thick PDMS slab to close the microchannel and leave an open chamber to allocate the villus-like microstructured hydrogel. The hydrogel was bonded using double-sided PSA rings. The microchannel and the bottom chamber were filled with PBS and prior to use, PBS was exchanged for the corresponding protein solution prepared in PBS-0.1% Tween20.

### In-chip diffusion studies on PEGDA-AA hydrogels using light-sheet fluorescence microscopy (LSFM)

A custom-made light-sheet microscope was developed for the visualization of the biomolecular gradients formed through the hydrogel network when allocated within the chip. Continuous wave lasers with wavelengths of 488 nm (MLD 50 mW, Cobolt) and 561 nm (DPL 100 mW, Cobolt) were used for fluorescence excitation. The light sheet was created by a pair of galvanometric mirrors (GVSM002, Thorlabs) conjugated with the illumination objectives (Plan-Fluor 4x, NA: 0.13, Nikon) through a pair of achromatic doublets (AC254-050-A-ML (f= 50 mm) and AC254-200-A-ML (f= 200 mm), Thorlabs), creating a flat top Gaussian beam profile to assure its homogeneity.^[66]^ The detection was performed in an up-right configuration, using water dipping objectives (PlanFluor 10x, NA: 0.3, Nikon) (Figure 2A, right). An achromatic doublet (AC254-200-A-ML, Thorlabs) formed an image onto a Hamamatsu Orca Flash 4.0 CMOS camera chip. Emission filters of 520/15 (for autofluorescence) and 620/52 (for Texas red) (Semrock) were selected using a motorized filter wheel (FW102C, Thorlabs). To be able to perform in-chip measurements, a special sample holder was designed using FreeCAD software and 3D printed (Hephestos 2, BQ) using poly(lactic acid) (PLA) as printing material (Figure 2A, right). It consisted of a square pool with top open access for the sample insertion and imaging, and two glass windows on the sides for proper light-sheet illumination. The chamber included a glass covered pedestal (oriented 25° from the optical table) where the microfluidic chip allocating the hydrogel was glued (Loctite). This pedestal also allowed bright-field illumination with a LED for sample positioning. With this inclination, the sample was illuminated and imaged transversally, to avoid any barrier on the path of the light-sheet. Laser scanning for imaging was performed by vertical translation of the sample holder using a motorized stage (PI M-501.1DG, Physik Instrumente), through a fixed horizontal light-sheet plane. All the components of the microscope were controlled using a custom-made software (LabView).

Biomolecular gradients were then formed by filling in the source compartment of the chip through the microfluidic channel through the inlet port while the hydrogel was in the allocated position. Biomolecular gradients formed by diffusing bovine serum albumin (BSA) protein fluorescently labelled with Texas Red (BSA_TxRED, Thermo Fisher) within the PEGDA-AA hydrogel networks were imaged by performing a time-lapse acquisition of a 3D volume of 1.33 × 1.33 × 2.3 mm over two hours with a frame interval of 15 min, and a stack step of 5 μm. The source chip compartment was loaded with BSA_TxRED dissolved in PBS-0.1% Tween20 at a concentration of 10 μg mL^-1^ and the imaging was started 3 hours after loading. To minimize data load, we binned the image by a factor of 4. Then, the overall lateral pixel size was 2.6 μm. The image processing was carried out using ImageJ software.^[67]^ First, the stacks were resliced (ImageJ reslice function) to obtain a vertical view of the source compartment, the hydrogel and the sink chip compartment. Then, a plane in the middle of the stack where the radial diffusion could be neglected was chosen. A linear region of interest (ROI) of 10 pixels in size was drawn along the perpendicular axis of the hydrogel. The relative fluorescence intensity (RFI) values along that line were obtained for all the time points acquired. These values were converted to protein concentrations by first constructing a calibration curve. For that, the fluorescence intensity of solutions with different BSA_TxRED concentrations (0 – 100 µg mL^-1^) were measured using the same LSFM set-up. For each concentration 21 measurements were made. Then, ROIs of 200 × 150 pixels in size were selected at the center of the stacks acquired in each measurement and the mean fluorescence intensity values were calculated. The background intensity value was determined from blank PBS.

### In-silico modelling of protein diffusion through PEGDA-AA hydrogels

In order to be able to estimate the spatio-temporal gradients of our molecules of interest (epidermal growth factor (EGF), R-Spondin 1, Noggin and Wnt2b) produced by the diffusion chip on the villus-like microstructured PEGDA-AA hydrogels, an in-silico model was developed. For that, finite element method (FEM) was employed. First, the diffusion coefficient of the proteins in an aqueous solution, D_0_, was calculated using Stokes-Einstein equation (**Equation 1**).

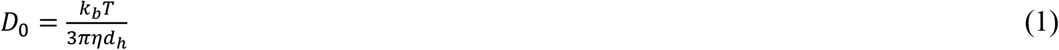

where k_b_ is the Boltzmann’s constant (1.38 × 10^−23^ J K^-1^), *T* is the absolute temperature (37°C = 310.15 K), *η* is the viscosity of the solvent (6.915 × 10^−4^ N m s^-2^ for water at 37°C^[68]^) and d_h_ is the protein hydrodynamic diameter. As d_h_ was not available in literature for our proteins of interest, for FEM simulations, the reported values of d_h_ corresponding to proteins with comparable sizes were selected (Table 1). The use of similar sized model proteins, whether for the simulations or experimental work, is accepted to provide a good estimate and it is a commonly adopted strategy in the literature for diffusing molecules that don’t interact with the hydrogel backbone (depending on the chemical composition of the hydrogel and the isoelectric point of the proteins).^[69,70]^

The diffusion of proteins over time and space was described by Fick’s second law and simulated employing the ‘Transport of diluted species’ module of COMSOL Multiphysics software (version 5.5). 3D forms were reduced to 2D geometries assuming axial symmetry. The geometry was composed of two media: the PEGDA-AA hydrogel and the aqueous solution. For each medium the corresponding diffusion coefficient values of each species (i.e. D_0_ for an aqueous solution, and D_g_ for a PEGDA-AA hydrogel network) used are listed in Table 2. D_0_ values were calculated using Stokes-Einstein equation and d_h_ values from Table 1. D_g_ values were assumed to be close to those reported in literature for model proteins diffusing through PEGDA-based hydrogels with similar network structures (PEGDA concentrations and chain length).

**Table 2:** For each species, diffusion coefficients values at 37°C in the aqueous solution (D_0_) and in the PEGDA hydrogel (D_g_); the source concentrations used.

| Model protein | $D_0 \cdot 10^{-10} [\text{m}^2 \cdot \text{s}^{-1}]$ | $D_g \cdot 10^{-10} [\text{m}^2 \cdot \text{s}^{-1}]$ | Concentration [$\text{ng} \cdot \text{mL}^{-1}$] | Molarity $\cdot 10^{-6} [\text{mol} \cdot \text{m}^{-3}]$ |
| --- | --- | --- | --- | --- |
| Insulin | 2.23 | 1.27 <sup>[68]</sup> | 300 | 52.6 |
| CA | 1.40 | 0.16 <sup>[68]</sup> | 600 | 20.7 |
| BSA | 0.91 | 0.13 <sup>[44]</sup> | 300 | 4.5 |
| Ovalbumin | 1.10 | 0.20 <sup>[68]</sup> | 600 | 14.0 |

As in the experimental setup, we modeled the gradients formation in a system composed by a source, the hydrogel, and a sink compartment. The initial concentration was set to zero in the entire geometry except for that of the source. The biomolecule concentrations commonly used in a standard organoid culture were tripled for the source (Table 2) to compensate for the biomolecules trapped at the membrane (≈ 66%), as determined in preliminary experiments (data not shown). Note that, even though the mass concentrations are similar or equal, the molarities are different due to differences in protein size. No flux boundary was imposed for the rest of the edges of the geometry. The villus-like microstructures were implemented in the simulation as an array of semi ellipses with a semi-minor axis of 75 μm (villus width) and a semi-major axis of 350 μm (villus height). The center-to-center spacing was set to 250 μm (corresponding to edge-edge spacing being 150 μm). The array was considered to be on top of a hydrogel base 150 μm thick (Figure 2B, left at the lower row). The sink medium was represented as a rectangular form. The FEM mesh for the aqueous medium was set to “finer” (10^−2^ mm < FEsize < 1 mm) and the mesh for the hydrogel medium was set to “extra fine” (10^−3^ mm < FEsize < 2·10^−2^ mm). To validate the simulations with the experimental data, BSA diffusion coefficients (in accordance with LSFM experiments) and a source concentration of 10 μg mL^-1^ were used. To fully capture experimental fluorescence profiles, a thin diffusion barrier between the source and the hydrogel was added to simulate the protein accumulation observed at the PET membrane (D = 5 × 10^−14^ m^2^ s^-1^). For the simulations emulating the gradient visualization experiments performed at RT using LSFM, D_0_ and D_g_ values for BSA at 25°C were used (D_0_ at 25°C = 6.8 × 10^−11^ m^2^ s^-1^ and D_g_ at 25°C = 1.0 × 10^−11^ m^2^ s^-1[44]^). This in-silico model was then used to predict the initial conditions of the spatio-temporal gradient profiles of the molecules of interest (EGF, R Spondin 1, Noggin, and Wnt2b) formed on the villus-like microstructured hydrogels. Note that, as the cell culture was performed in a Transwell culture setup (see ahead), the source and the sink dimensions were adjusted accordingly. The diffusion coefficients and source concentrations detailed in Table 2 were employed for the simulations. The simulations were performed up to 24 h of diffusion time, which was found to be a convenient frequency for the cell culture medium exchange and also it was confirmed with our simulations that gradients profiles could be stabilized by this periodic medium replenishment approach (see Supporting Information for details). As results, the factor concentration profiles as a function of the vertical position (z) along a line spanning the source, the hydrogel and the sink (Figure 2B, bottom left), and also along the curve contouring the villus-like surface were computed at different time intervals (see the corresponding figure legends) and the graphs were plotted using GraphPad software. Base to tip decrease in biomolecular concentrations was calculated using the concentration values computed at the base and the tip for each factor.

### Mouse model

Lgr5-EGFP-IRES-creERT2 mice, previously described,[3] were used. Briefly, Lgr5-EGFP-IRES-creERT2 mice were generated by homologous recombination in embryonic stem cells targeting the EGFP-IRES-creERT2 cassette to the ATG codon of the stem cell marker Lgr5 locus, allowing the visualization of Lgr5^+^ stem cells with a green fluorescent protein (GFP). All experimental protocols involving mice were approved by the Animal care and Use Committee of Barcelona Science Park (CEEA-PCB) and the Catalan government and performed in accordance with their relevant guidelines and regulations.

### Intestinal crypt isolation and culture

Mice intestinal crypts were isolated as previously described.^[71,72]^ In short, small intestines were washed with PBS and cut longitudinally. Villi were mechanically removed, and intestinal crypts were extracted by incubating the tissue with 2 mM of EDTA (Sigma-Aldrich) in PBS for 40 minutes at 4°C. The digestion content was filtered through a series of 70 µm pore cell strainers (Biologix Research Co.) to obtain the crypt fractions. Crypts were resuspended in Matrigel (BD Bioscience) drops and cultured with basic organoid growth medium (Advanced DMEM/F12 (Invitrogen), Glutamax (1%, Gibco), HEPES (1%, Sigma-Aldrich), Normocin (1:500, Invitrogen), B27 (2%, Gibco), N2 (1%, Gibco), N-acetylcysteine (1.25 mM, Sigma-Aldrich)), supplemented with recombinant murine EGF (100 ng mL^-1^, Gibco), recombinant human R-spondin 1 (200 ng mL^-1^, R&D Biosystems), recombinant murine Noggin (100 ng mL^-1^, Peprotech), CHIR99021 (3 μM, Tebu-bio) and valproic acid (1 mM, Sigma-Aldrich) to obtain ENRCV medium. The first 4 days of culture the Rho kinase inhibitor Y-27632 (Sigma-Aldrich) was added. The medium was changed every 2 to 3 days. Outgrowing crypts were passaged twice a week and organoid stocks were maintained for up to 4 months.

### Intestinal subepithelial myofibroblast conditioned medium (ISEMF_CM) preparation

Intestinal subepithelial myofibroblasts (ISEMFs) were isolated from mice intestines following a previously reported protocol.^[73]^ Briefly, left over tissue from the crypt isolation procedure was further digested with 2000 U of collagenase (Sigma-Aldrich) for 30 minutes at 37°C. The digested tissue was centrifuged and the pellet was resuspended in DMEM, high glucose, GlutaMAX ™, pyruvate (Life Technologies) containing fetal bovine serum (FBS) (10%, Gibco), penicillin/streptomycin (1%, Sigma-Aldrich), and minimal essential medium non-essential amino acids (MEM-NEAA) (1%, Gibco) and cultured in tissue culture plates. After 1 week in culture, lamina propria myofibroblasts positive for α-smooth muscle actin (α-SMA) and vimentin and negative for desmin (data not shown) remained attached.^[10]^ Plates at 60 to 80% of confluence were maintained in culture for 6 days and then the supernatant was collected and complemented with B27 (2%), N2 (1%) and N-acetylcysteine (1.25 mM) to obtain what we called intestinal subepithelial myofibroblast conditioned medium (ISEMF_CM).

### Formation of biomolecular gradients along the vertical axis of the villus-like microstructured PEGDA-AA hydrogels

Gradients of the biomolecules reported to be relevant for the formation and maintenance of the intestinal stem cell (ISC) niche at the intestinal crypts, listed in Table 1, were established as predicted by the in-silico model. For that purpose, villus-like microstructured hydrogels fabricated on top of PET membranes were mounted on Transwell inserts, which allow the use of different solutions at the apical and basolateral compartments. To account for the cellular effects of the biomolecular gradients formed, four different culture conditions were established: (i) asymmetric, where ENRCV and ISEMF_CM were delivered basolaterally and basal medium was delivered apically; (ii) uniform, with the basolateral and apical delivery of ENRCV and ISEMF_CM; (iii) asymmetric 2.0, where the concentrations of Noggin and R-spondin were doubled compared to asymmetric while keeping basal medium at the apical chamber, and (iv) asymmetric Wnt2b, where the ISEMF_CM was replaced by the recombinant protein Wnt2b, so ENRCV and Wnt2b were delivered in the basolateral chamber and basal medium at the apical side. Final concentrations for the different relevant biomolecules (Table 2) were determined from the simulations and are detailed at Table S1. Notice that, in order to compensate for ≈ 66% of protein concentration getting trapped at the membrane, protein source concentrations were tripled compared to concentrations used for organoids culture within Matrigel drops. For all four conditions, the different media were introduced in the corresponding Transwell compartment 24 hours before cell seeding and, according to the simulation data, we renewed the medium of both compartments every 24 hours to maintain the gradients formed.

### Organoid-derived cell culture onto villus-like PEGDA-AA microstructured hydrogels with biomolecular gradients

Matrigel drops containing fully-grown organoids were shattered by pipetting with TrypLE Express1X (Gibco) and passed several times through a syringe with a 23 G needle (BD Microlance 3). Next, disrupted organoids were further digested to single cells by incubating them for 5 to 7 minutes at 37°C with vigorous hand-shaking every minute. Successful digestion to single cells was confirmed via inspection under the microscope. Prior to cell seeding on the villus-like microstructured hydrogels, the media of both apical and basolateral compartments of the Transwells were renewed to maintain the established gradients. Then, 5×10^5^ intestinal epithelial cells (the cell number was optimized to obtain full surface coverage on day 3 and adjusted slightly for each crypt isolation) were resuspended in 100 µL of the corresponding apical medium and seeded on the hydrogels. Once the cells were adhered (after 1 hour), 100 µL of the corresponding apical medium were added. Cell cultures were maintained in an incubator at 37°C and 5% CO_2_ for at least one day.

### Immunostaining

Cells were fixed with 10% neutralized formalin (Sigma-Aldrich), permeabilized with 0.5% Triton X-100 (Sigma-Aldrich), and blocked with a blocking buffer containing 1% BSA (Sigma-Aldrich), 3% donkey serum (Millipore), and 0.2% Triton X-100 in PBS for at least 2 hours at room temperature (RT). The primary antibodies used were: anti-Ki67 (1:100, Abcam), anti-Ki67 (1:100, BD Biosciences), anti-GFP (1:100, Abcam), anti-Lysozyme (Lyz) (1:100, Dako), anti-Fatty acid binding protein 1 (FABP1) (1:50, Sigma-Aldrich) and anti-cytokeratin 20 (CK20) (1:100, Dako). All samples were incubated with the primary antibody overnight at 4°C followed by 1 h incubation at RT with secondary antibodies, Alexa Fluor 568 and Alexa Fluor 488 donkey anti-goat, and Alexa Fluor 647 (Jackson ImmunoResearch) diluted at 1:500. Nuclei were stained with 4’,6-diamidino-2-phenylindole (DAPI) (1:1000, Thermo Fisher Scientific). Alexa Fluor 568 phalloidin was used to stain filamentous actin (F-actin) (3.5:500, Cytoskeleton). For detection of mucus secreting goblet cells, periodic acid-Schiff (PAS) (Sigma-Aldrich) was used.

### Image acquisition and analysis

Fluorescence images were acquired at randomly selected locations using a confocal laser scanning microscope (LSM 800, Zeiss) with a 10x (NA = 0.3, WD = 2.0) and a confocal spinning disk microscope (LIPSI, Nikon) with a 20x air objectives (NA = 0.7, WD = 2.3). The laser excitation and emission light spectral collection were optimized for each fluorophore, especially for the four-color scans, where the emission bands were carefully adjusted to avoid overlapping channels. The pinhole diameter was set to 1 Airy Unit (AU). A z-step of 5 µm was used. The relative coverage was quantified by measuring the cell area after thresholding with ImageJ software the maximum intensity projected F-actin fluorescence images and normalizing to surface of the substrate (base + pillars). For this, we employed a model to estimate the effective surface of the substrate from the projected image (see Supporting Information for the details). In addition, confocal fluorescence microscopy images of Ki67, GFP, Lysozyme, CK20 and FABP1 markers were used to quantify the proliferative cells, stem cells, Paneth cells, differentiated cells and enterocytes, respectively, along the vertical axis of the villus-like microstructures. For this, a custom-made ImageJ macro was used to divide the z confocal stacks to tiles, each comprising one single villus. For Ki67 and FABP1 markers, we used the following procedure. Each covered pillar was analyzed using IMARIS (BitPlane, version 9.1.0). As a pre-processing step, we applied a mean filter of 3 × 3 × 3 px to the Ki67 channel to reduce the dotty unspecific staining of this antibody. Then, nuclei were detected with the Imaris Spot detector (diameter set to 9 μm) by adjusting the threshold manually for optimal results. Next, positive cells for the marker in question (Ki67 or FABP1) were identified by measuring the mean intensity on the positions of the detected nuclei and setting a minimum threshold for positive cells. Finally, we exported the Z positions of all cells and positive cells. For each pillar, Z values were rescaled from 0 to 1, 0 corresponding to the nucleus detected closer to the villus base and 1 to the nucleus closer to the villus tips. Z was divided into five equal segments for data interpretation purposes. Next, we estimated the percentage of positive cells with respect to the total number of cells for the five regular Z segments along each villus. Finally, we computed the fractioned percentage of each segment. To quantify the relative change in the frequency of Ki67^+^ cells from day one to day three, we subtracted the mean frequency of Ki67^+^ cells of day 1 to that of day 3 and we normalized the value by the mean frequency of day 1. To quantify the intensity signal along the vertical axis of each villus of CK20, GFP and Lyz, we used another custom-made ImageJ macro. Briefly, after dividing each pillar into individual stacks, we run the ImageJ built-in plot profile function within a drawn ROI surrounding the villus to quantify mean intensity for the marker in question (CK20, GFP or Lyz) and F-acting fluorescent channels and we exported the data. We normalized the intensity values of the marker of interest by the F-actin ones and we rescaled them from 0 to 1 to eliminate the intensity variations due to the image acquisition set up. Then, we rescaled the villus vertical axis from 0 to 1 and divided it into five equal segments for data interpretation purposes. Finally, we computed the mean intensity fraction of each segment.

### Statistical analysis

N is the number of independent experiments, n is the number of samples and the word pillars indicates the number of pillars analyzed. The data were presented in the figures as mean ± error (standard error of the mean (SEM) or standard deviation (SD) as indicated in the figure captions). Statistical comparisons were performed using two-tailed, unequal variance t-test, an alpha of 0.05 and a p < 0.05 (* < 0.05, ** < 0.01 and *** < 0.001) was considered to be significant. The software used for statistical analysis was GraphPad Prism (GraphPad software, La Jolla, CA, USA).

## Supporting Information

Supporting Information is available from the Wiley Online Library or from the author.

## Acknowledgements

We would like to thank Rene Fabregas for the advice on the in-silico model. We also thank Jordi Comelles, María García and Enara Larrañaga for the fruitful discussions. Funding for this project was provided by European Union’s Horizon 2020 ERC grant agreements No 647863 (COMIET) and No 884623 (residualCRC), the CERCA Programme/Generalitat de Catalunya (2017-SGR-1079 and 2017-SGR-698), Laserlab-Europe EU-H2020 grant agreement No 871124, the Spanish Ministry of Economy and Competitiveness (Severo Ochoa Program for Centers of Excellence in R&D 2016–2019 and CEX2019-000910-S), the Spanish Ministry of Science (PID2020-119917RB-I00) and also by the Fundació Privada Cellex and the Fundació Mir-Puig. The authors would like to thank Barcelona Institute of Science and Technology (BIST) for the funding of ENGUT project through Ignite Programme. The authors gratefully acknowledge the Agència de Gestió d’Ajuts Universitaris i de Recerca (AGAUR) for the funding of G. A. through FI-DGR 2014 and to MINECO/FEDER Ramón y Cajal program (RYC-2015-17935) for the funding of E. G. The results presented here reflect only the views of the authors; the European Commission is not responsible for any use that may be made of the information it contains.

Received: ((will be filled in by the editorial staff)) Revised: ((will be filled in by the editorial staff)) Published online: ((will be filled in by the editorial staff))

