## Supporting information for "Modelling biochemical gradients in vitro to control cell compartmentalization in a microengineered 3D model of the intestinal epithelium"

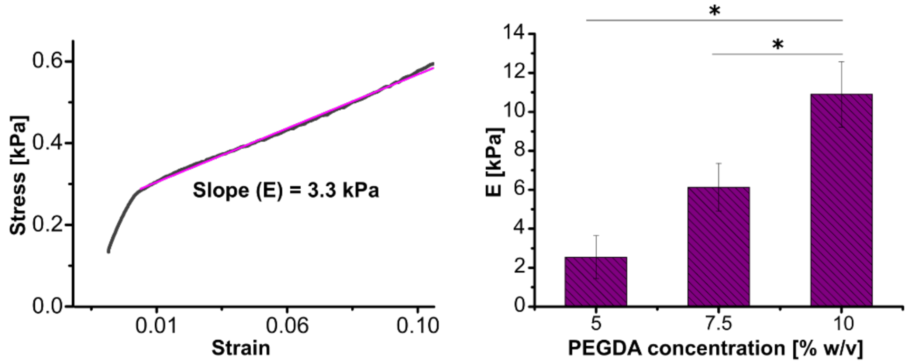


**Figure S1.** Stress-strain curve of a 5% w/v PEGDA-AA hydrogel obtained by a compression test (left). The Young's modulus (E) was calculated from the slope of the linear part of the stress-strain curves. Graph showing the Young's modulus of PEGDA-AA hydrogels at different concentrations measured after swelling (n=3) (right). P-value: *p<0.05. Mean ± SD. The Young's modulus values determined varied from 2.5 ± 1.1 kPa to 10.9 ± 1.7 kPa within 5% - 10% w/v PEGDA concentration range, consistent with the values reported for small intestine.[1]

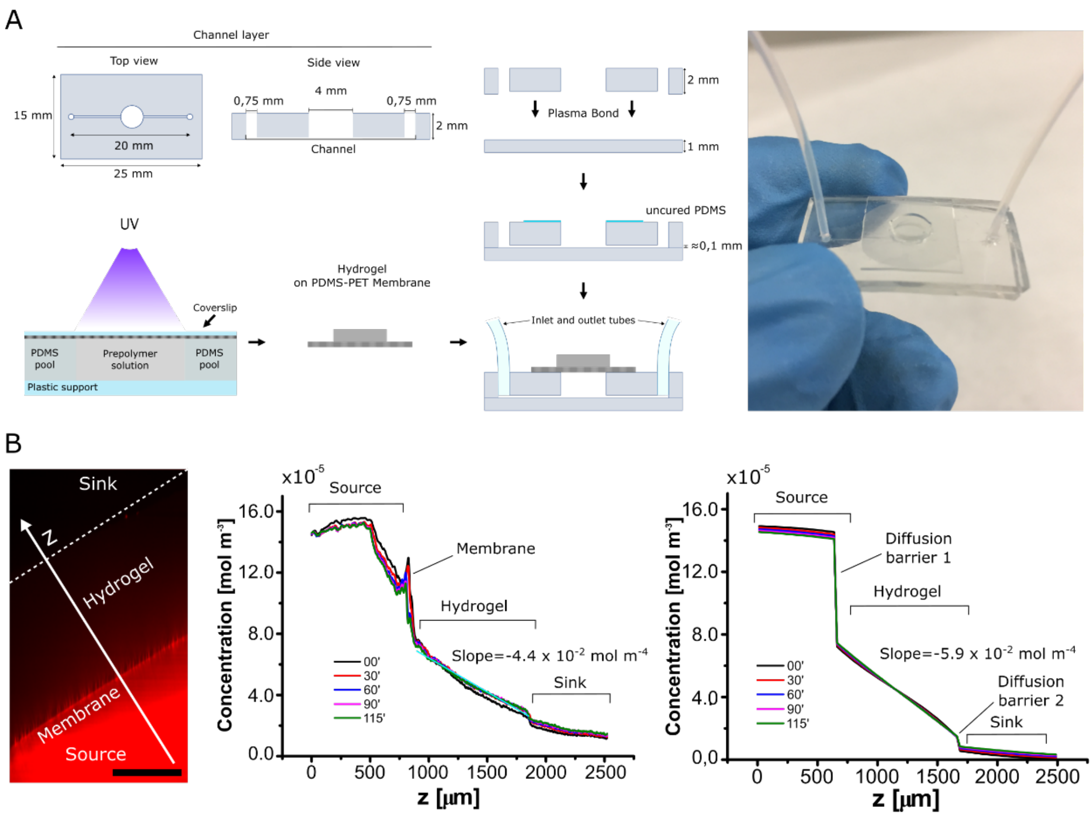


**Figure S2.** (A) Schematic showing the preparation together with the dimensions of the PDMS layer with the channel (104.3 ± 0.1 µm) and the bonding of the hydrogels to the chip with a thin layer of uncured PDMS (left). Image showing the assembled chip allocating the hydrogel disc (right). (B) Cross-sectional fluorescence image acquired by LSFM spanning the source, the hydrogel and the sink of Texas Red-labeled BSA (red) diffusion through hydrogel discs at 24 h after loading (left). Scale bar: 500 µm. Graph showing the concentration profiles taken along the white arrow shown in the fluorescence image on the left at different times points (middle). Concentration profiles predicted by the FEM model at the corresponding time points (right). The beginning of the time-lapse acquisition was taken as the zero time point.

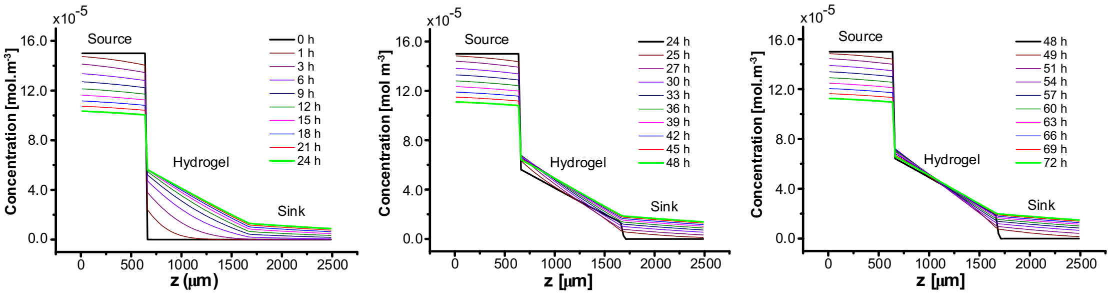

**Figure S3.** Graphs showing the concentration profiles taken along a perpendicular line spanning the source, the hydrogel and the sink in FEM simulation ran up to 72 h with medium change simulation implemented at 24 h and 48 h. Hydrogel disc geometry and model protein BSA were used for the simulations.

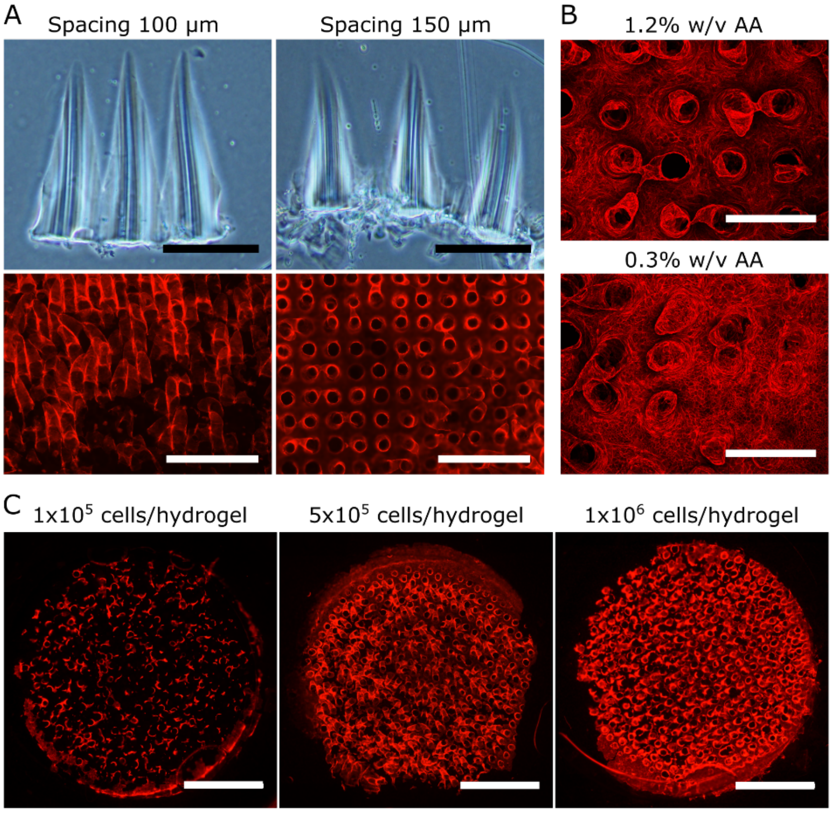


**Figure S4.** (A) Bright field microscope images of the cross-section of the villus-like microstructured hydrogel scaffolds fabricated with a spacing of 100 mm (top left) and 150 mm (top right). The hydrogel formulation used for the base formation was: 6.5% w/v PEGDA and 1.2% w/v AA. The epithelial cells were cultured in the presence of the gradients. Scale bars: 400 mm. Representative wide field images showing fluorescence for F-actin of epithelial cells cultured for 4 days on scaffolds fabricated with 100 mm (bottom left) and 150 mm (bottom right). Scale bars: 1 mm. (B) Representative confocal maximum projection images showing fluorescence for F-actin of epithelial cells cultured for 4 days on scaffolds fabricated with 1.2% w/v AA (top), and 0.3% w/v AA (bottom) in the hydrogel formulations used for the base formation. The PEGDA concentration used was kept the same i.e. 6.5% w/v for both the pillar and the base formation. The epithelial cells were cultured in the presence of the gradients. Scale bars: 400 µm. (C) Representative wide field tile scan images showing fluorescence for F-actin of epithelial cells seeded at $1x10^5$ cells/sample (left), $5x10^5$ cells/sample (middle), and $1x10^6$

cells/sample (right) seeding densities and cultured for 3 days on scaffolds fabricated with 150 spacing and 6.5% w/v PEGDA and 0.3% w/v AA hydrogel formulation for both the pillar and the base formation. The tile scan images show the entire Transwell membrane surface (0.33 $cm^2$). The epithelial cells were cultured in the presence of the gradients. Scale bars: 2 mm.

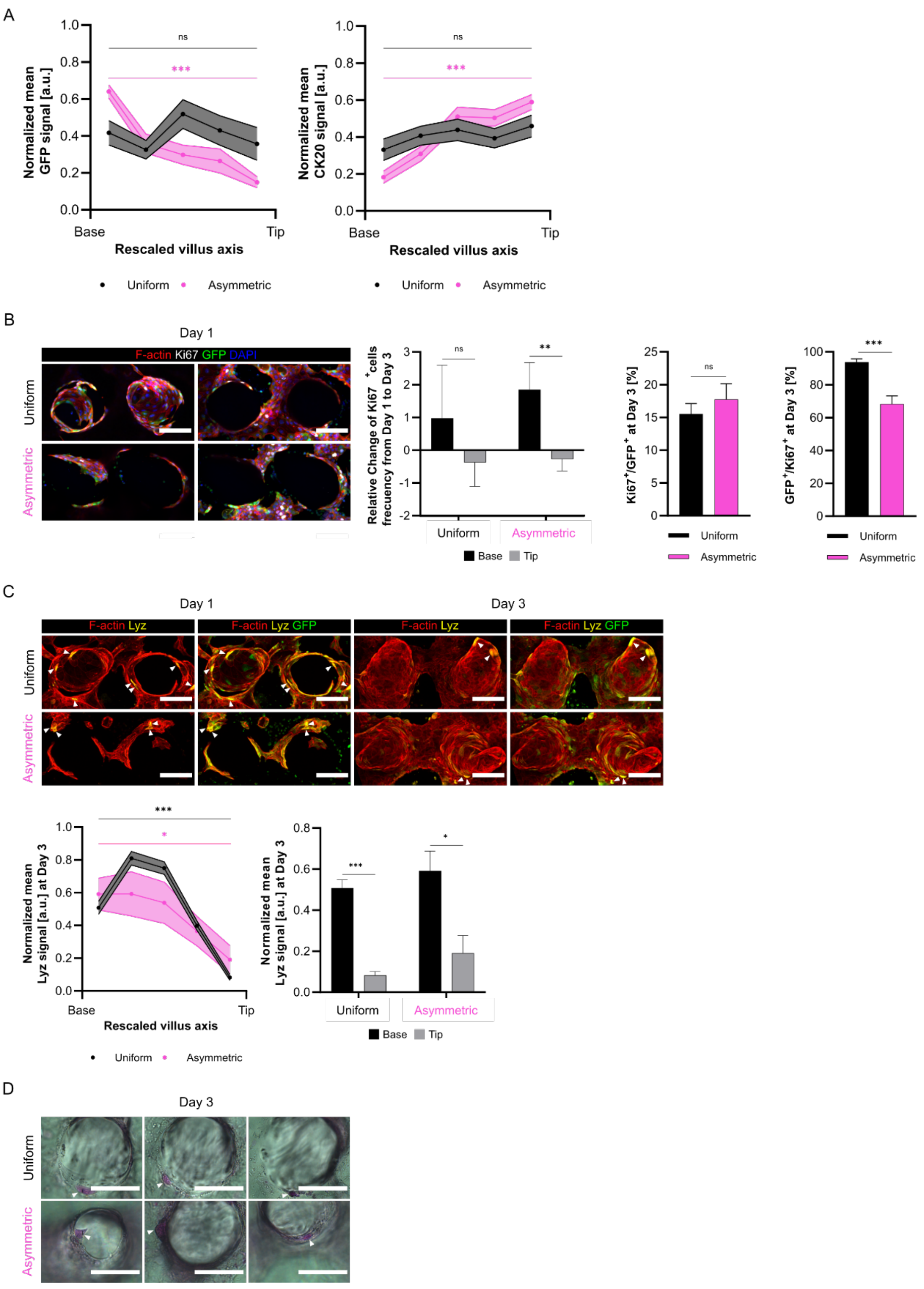


**Figure S5.** (A) Normalized mean GFP signal along the rescaled villus axis in Asymmetric or Uniform conditions at day 3 of culture (right). Uniform: N = 1, pillars = 12. Asymmetric: N = 1, pillars = 12. P-value: *** < 0.001. Mean ± SEM. Normalized mean CK20 signal along the rescaled villus axis in Asymmetric or Uniform conditions at day 3 of culture (right). Uniform: N = 2, pillars = 19. Asymmetric: N = 4, pillars = 41. P-value: *** < 0.001. Mean ± SEM. (B) Representative confocal images of Uniform (top) and Asymmetric (bottom) samples on day 1 of culture immunostained for F-actin, Ki67, GFP and DAPI, Maximum intensity projection of the section corresponding to the pillar (left) and to the base (right). Scale bars: 100 μm. Frequency of $GFP^+$ cells within $Ki67^+$ population at day 3. Uniform: N = 1, pillars = 19. Asymmetric: N = 1, pillars = 10. P-value: *** < 0.01. Mean ± SEM. Relative change of $Ki67^+$ cells from Day 1 to Day 3 at the base and pillar sections for Uniform and Asymmetric (middle). Uniform: Day 1 N = 1, pillars = 3, day 3 N = 2, pillars = 19. Asymmetric: Day 1 N = 2, pillars = 19, day 3 N = 4, pillars = 41. P-value: ** < 0.01. Mean ± SD. (C) Confocal images of samples on day 1 (top) and day 3 (bottom) of culture of Uniform and Asymmetric conditions immunostained for F-actin, Lyz (Paneth cells) and GFP. Maximum intensity projections. Arrow heads point to $Lyz^+$ cells. Scale bars: 100 μm. Normalized mean Lyz signal along the rescaled villus axis and at the base and the tip of the pillars in Asymmetric or Uniform conditions at day 3 of culture (right). Uniform: N = 1, pillars = 12. Asymmetric: N = 1, pillars = 12. P-value: * < 0.05, *** < 0.001. Mean ± SEM. (D) Bright field images showing $PAS^+$ positive Goblet cells marked with arrow heads present in Uniform (top) and Asymmetric (bottom) conditions on day 3. Scale bars: 100 μm.

**Characterization of the mesh size of PEGDA-AA hydrogels by swelling**

The mesh size of the PEGDA-AA hydrogels was estimated using Flory-Rehner equilibrium swelling theory as modified by Peppas and Merrill for hydrogels diluted in aqueous media.[2–4] (1) First, the measured $m_c$, $m_s$, and $m_d$ values of PEGDA-AA hydrogels (see Materials and Methods) were used to calculate weight fractions of the hydrogels after fabrication ($q_F$) and at equilibrium swelling ($q_W$) by the following equations

$$q_F = \frac{m_c}{m_d} \tag{1}$$

$$q_W = \frac{m_s}{m_d} \tag{2}$$

Then, the polymer volume fraction of the hydrogel in swollen state ($v_{2,s}$) and the volume fraction in the relaxed state ($v_{2,r}$) were calculated, by the equations (3) and (4), respectively.

$$v_{2,s} \left[1 + \frac{(q_W - 1) r_p}{r_{swel}}\right]^{-1} \tag{3}$$

$$v_{2,r} = \left[1 + \frac{(q_F - 1) r_p}{r_{sol}}\right]^{-1} \tag{4}$$

where ρ is the density the PEG (1.12 g cm$^{-3}$),[5,6] $\rho_{swel}$ is the density of swelling agent and $\rho_{sol}$ is the density of the solvent in which the PEGDA solution was prepared, both of which was PBS in our setup. The density of PBS was taken as 1.01 g cm$^{-3}$ (calculated from the product datasheet).[3,5,7] Then, the molecular weight between crosslinks ($M_c$)[2–4] was computed using with the equation below (5).

$$\frac{1}{M_c} = \frac{2}{M_n} - \frac{\frac{v}{V_1}\left[\ln(1 - v_{2,s}) + v_{2,s} + \chi v_{2,s}^2\right]}{v_{2,r}\left[\left(\frac{v_{2,s}}{v_{2,r}}\right)^{1/3} - \frac{1 v_{2,s}}{2 v_{2,r}}\right]} \tag{5}$$

where $M_n$ is the average molecular weight of the polymer, which is 6000 Da (P6000) in our case, v is the specific volume of bulk PEGDA (0.893 cm$^3$ g$^{-1}$),[8] $V_1$ is the molar volume of the solvent which is approximated to the molar volume of water (18 cm$^3$ mol$^{-1}$)[8] and finally, χ is the value of the polymer-solvent interaction parameter (0.426 as determined by Merill *et al.*).[9] For the determination of the mesh size (ξ), the equations described by Canal and Peppas were used.[10] For that, first, the root mean square of the end-to-end distance of the unperturbed state of the PEGDA chain was calculated using the following equation (6).

$$(\bar{r}_0^{2})^{1/2} = l\, C_n^{1/2}\, n^{1/2} \tag{6}$$

where $C_n$, the characteristic ratio, that is 4,[9] and l, the weighted average bond length of the PEG backbone, is 0.15 nm.[8] The n is the number of repeating units in the crosslink, which depends on $M_c$ and $M_r$, the molecular weight of the PEG repeating unit (44 g mol$^{-1}$).[8] The n was determined by the equation below (7).

$$n = 2\frac{M_c}{M_r} \tag{7}$$

Finally, the mesh size (ξ) was calculated by using the equation (8).

$$\xi = v_{2,s}^{-1/3}\, (\bar{r}_0^{2})^{1/2} \tag{8}$$

Four different hydrogel samples were analyzed for each condition (n=4). MATLAB software was used to compute the values and the data was plotted as mean ± standard deviation with GraphPad software. The statistical comparison was performed using two tailed, unequal variances t-test and $p<0.05$ was considered significant.

**Characterization of the mechanical properties of PEGDA-AA hydrogels**

Mechanical properties of the hydrogels were analyzed by a compression test. The gels were casted as discs, 10 mm in diameter and ≈ 1 mm in height, and swollen to constant weight. The concentrations characterized were: 5% w/v, 7.5% w/v, and 10% w/v P6000 prepared in PBS with 1% w/v I2959 and 0.06% w/v AA (mechanical properties do not change significantly from 0.06% w/v to 1.2% w/v AA[11]). The UV exposure time used for the fabrication of the hydrogels was 200 s. Stress-strain curves were obtained in compression mode with a dynamic mechanical analyzer equipment (Q800 Dynamic Mechanical Analyzer, TA Instruments). The samples were placed in between the compression clamps and deformed by applying a constant strain rate of $1.20\% \cdot min^{-1}$ under unconfined compression up to a maximum strain of 10%, in agreement with similar works reported in the literature.[12,13] A small preload of 100 mN was used to promote an adequate contact between the hydrogels and the apparatus. The Poisson ratio was selected to be 0.5, in accordance with previously published reports.[12,13] The cross-sectional area, A, of the hydrogels and the Poisson ratio were introduced to the operational software of the equipment (TA Instruments). The stress, $\sigma$, was calculated by the normal force applied, F, divided by A; and the strain, $\varepsilon$, as the ratio of the change in the hydrogel height to the original height. Then, the stress-strain curves were plotted and from the slope of the linear part of the curves, Young's modulus (E) was calculated. At least three hydrogel samples were analyzed for each condition. The data were plotted as mean ± standard deviation with GraphPad software. The statistical comparison was performed using two tailed, unequal variances t-test and $p<0.05$ was considered significant.

**In-silico modelling of protein diffusion on PEGDA-AA hydrogels**

To simulate the effect of the medium change cycles (performed experimentally every 24h) on the gradient profiles a simpler rectangle model was used. For that, sequential runs were performed using the hydrogel gradient profiles from the previous last time point as the initial condition for the hydrogel in the following simulation; while resetting the source and the sink to their initial values. The aim of doing this was to simulate the effect of the medium change on the gradient profiles.

**Formation of ISC niche biomolecule gradients through the hydrogels**

**Table S1:** For each regime and each compartment, the medium used and the concentrations of each species.

| Asymmetric [illegible] | Asymmetric 2.0 | | Asymmetric | | Uniform | |
|---|---|---|---|---|---|---|
| **Bottom** | **Top** | **Bottom** | **Top** | **Bottom** | **Top** | **Bottom** |
| **Basal medium Wnt2b 3 ($EN_{2x}R_{2x}CV$)** | **Basal medium** | **ISEMF_CM 3· ($EN_{2x}R_{2x}CV$)** | **Basal medium** | **ISEMF_CM 3 (ENRCV)** | **ISEMF_CM 1 (ENRCV)** | **ISEMF_CM 3 (ENRCV)** |
| 300 | 0 | 300 | 0 | 300 | 100 | 300 |
| 600 | 0 | 600 | 0 | 300 | 100 | 300 |
| 1200 | 0 | 1200 | 0 | 600 | 200 | 600 |
| 9 | 0 | 9 | 0 | 9 | 3 | 9 |
| 3 | 0 | 3 | 0 | 3 | 1 | 3 |
| 600 | 0 | 0 | 0 | 0 | 0 | 0 |

| | COMPARTMENT | MEDIUM | EGF (ng mL$^{-1}$) | Noggin (ng mL$^{-1}$) | R-Spondin1 (ng mL$^{-1}$) | CHIR (µM) | Valproic Acid (mM) | Wnt2b (ng mL$^{-1}$) |
|---|---|---|---|---|---|---|---|---|
| | Top | Basal medium | 0 | 0 | 0 | 0 | 0 | 0 |

**Surface Coverage Quantification**

3D surface coverage was quantified using the F-actin fluorescence images. First, the pillar area was calculated by approximating it to that of a truncated cone (ATC). Next, the total number of pillars in a given area was estimated. For that, first the area occupied by 4 pillars was calculated by selecting a square ROI. Then, this number was proportioned to the 2D projected total area of the villus-like hydrogel to obtain the total number of pillars (p).

The true area (TA) can then be calculated as follows:

$$TA = 2D_{projected\ total\ area} - Base_{covered\ area} + Pillars_{covered\ area}$$

$$TA = 2D_{projected\ total\ area} - p * (\pi R^2) + p * ATC$$

The truly uncovered area (TUA) was calculated as follows:

$$TUA = 2D_{projected\ uncovered\ area} - Base_{unarea} + Pillars_{uncovered\ area}$$

$$TUA = 2D_{projected\ uncovered\ area} - p_{uncovered} * (\pi R^2) + p_{uncovered} * ATC$$

To quantify the $2D_{projected\ uncovered\ area}$, the $2D_{truly\ covered\ area}$, measured using the threshold function in ImageJ, was subtracted to the $2D_{projected\ total\ area}$ . Finally, surface coverage % was calculated as follows:

$$Surface\ Coverage\ (\%) = \frac{TA - TUA}{TA} * 100$$


[1] J. Sotres, S. Jankovskaja, K. Wannerberger, T. Arnebrant, *Sci. Rep.* **2017**, *7*, 1.

[2] P. J. Flory, J. Rehner, *J. Chem. Phys.* **1943**, *11*, 521.

[3] N. A. Peppas, E. W. Merrill, *J. Appl. Polym. Sci.* **1977**, *21*, 1763.

[4] N. A. Peppas, P. Bures, W. Leobandung, H. Ichikawa, *Eur. J. Pharm. Biopharm.*

**2000**, *50*, 27.
